# Development of a functional genetic tool for Anopheles gambiae oenocyte characterisation: application to cuticular hydrocarbon synthesis

**DOI:** 10.1101/742619

**Authors:** Amy Lynd, Vasileia Balabanidou, Rudi Grosman, James Maas, Lu-Yun Lian, John Vontas, Gareth J. Lycett

**Affiliations:** Vector Biology Dept, Liverpool School of Tropical Medicine, Liverpool, L3 5QA, UK; Institute of Molecular Biology and Biotechnology, Foundation for Research and Technology-Hellas, Heraklion 70013, Greece; Institute of Integrative Biology, University of Liverpool, Liverpool, UK; Department of Crop Science, Agricultural University of Athens, 11855 Athens, Greece

**Author notes:** Corresponding Author Dr. Gareth Lycett, Vector Biology Dept., Liverpool School of Tropical Medicine, Pembroke Place, Liverpool, L3 5QA, UK.

**Keywords:** cytochrome P450, gene knockdown, malaria, enhancer, Gal4 driver

## Abstract

Oenocytes are an insect cell type having diverse physiological functions ranging from cuticular hydrocarbon (CHC) production to insecticide detoxification that may impact their capacity to transmit pathogens. To develop functional genetic tools to study *Anopheles gambiae* oenocytes, we have trapped an oenocyte enhancer to create a transgenic mosquito Gal4 driver line that mediates tissue-specific expression. After crossing with UAS-reporter lines, *An. gambiae* oenocytes are fluorescently tagged through all life stages and demonstrate clearly the two characteristic oenocyte cell-types arising during development. The driver was then used to characterise the function of two oenocyte expressed *An. gambiae cyp4g* genes through tissue-specific expression of UAS-RNAi constructs. Silencing of *cyp4g16* or *cyp4g17* caused lethality in pupae of differing timing and penetrance. Surviving *cyp4g16* knockdown adults showed increased sensitivity to desiccation. Total cuticular hydrocarbon levels were reduced by approximately 80% or 50% in both single gene knockdowns when assayed in young pupa or surviving adults respectively, indicating both genes are required for complete CHC production in *An. gambiae* and demonstrate synergistic activity in young pupae. Comparative CHC profiles were very similar for the two knockdowns, indicating overlapping substrate specificities of the two enzymes. Differences were observed for example with reduced abundance of shorter chain CHCs in CYP4G16 knockdowns, and reduction in longer, branched chained CHCs in CYP4G17 knockdown adults. This is the first time that two *cyp4g*s have both been shown to be required for complete CHC production in an insect. Moreover, the generation of tagged cells and identification of an enhancer region can expediate oenocyte specific transcriptomics. The novel driver line can also be used to explore oenocyte roles in pheromone production, mating behaviour and longevity in the malaria mosquito.

## Introduction

Oenocytes are insect secretory cells that in diptera such as Drosophila exist as morphologically distinct derivatives in larvae and adults with separate developmental origins (Gould et al. 2001). In the fruitfly, larval oenocytes originate from embryonic ectodermal cells, whereas adult oenocytes are thought to derive from pupal histoblasts (Makki et al. 2014). These liver-like cells that have diverse physiological functions encompassing regulation of respiration, tissue histolysis, detoxification, hormone production, dietary related longevity and cuticle synthesis (Makki et al. 2014; Stefana et al. 2017). Perhaps best studied to date are those roles linked to lipid metabolism. Fruitfly oenocytes express an abundance of lipid synthesis enzymes, including those in the pathways leading to the production of very long chain (VLC) lipids and cuticular hydrocarbons (CHC) (Gutierrez et al. 2007). These are transported to the tracheal system and outer insect integument to provide a hydrophobic waxy layer to maintain water balance (Parvy et al. 2012). This key adaptive trait provides insects with the ability to limit dehydration, yet repel water to prevent flooding, in order to occupy different niches. The precise control of CHC composition to maintain water balance is compounded in insects with aquatic immature stages, including mosquitoes, which must switch rapidly during pupal development from submersion in water to an environment with variable humidity.

Our interest in developing genetic tools to study mosquito oenocytes stems from an initial observation that the sole reductive binding partner, cytochrome p450 reductase (CPR), of the largest family of multifunctional oxidative enzymes, the cytochromes P450 (CYPs), was massively expressed in these cells, in both *An. gambiae* and Drosophila (Lycett et al. 2006). Later studies in Drosophila linked expression of CPR and a single CYP4G partner, CYP4G1 in oenocytes with the final decarbonylation step in the cuticular hydrocarbon synthetic pathway that converts VLC aldehydes to alkanes and alkenes (Qiu et al. 2012a). Similar activity has recently been detected in the single *cyp4g* gene encoded in the honey bee genome (Calla et al. 2018). In contrast two *cyp4g* genes are annotated in the Anopheles genome, *cyp4g16* and *cyp4g17*, and we have previously shown that both gene products are highly expressed in adult and pupal oenocytes (Kefi et al. 2019; Ingham et al. 2014; Balabanidou et al. 2016), yet only CYP4G16 had decarbonylase activity *in vitro*. In other species with two annotated members of the *cyp4*g family, both recombinant enzymes possess decarbonylase activity (MacLean et al. 2018), and when the Anopheles genes were expressed in Drosophila in a *cyp4g1* silenced background either enzyme or in combination rescued the lethal phenotype (Kefi et al. 2019). The observed inactivity of the recombinant *An. gambiae* CYP4G17 is thus most likely due to incompatibility with the low molecular weight substrate aldehyde that was tested *in vitro* (Balabanidou et al. 2016).

To expand the *in vivo* analysis of oenocyte function in mosquitoes, we have developed a Gal4 driver line to modify gene expression specifically in *An. gambiae* oenocytes, and have initially targeted the *cyp4g* genes. Our earlier research identified a number of Gal4 activators and UAS modules suitable for expression in *An. gambiae* (Lynd and Lycett 2012, 2011). Moreover, in creating a series of docking lines for recombinase mediated cassette exchange (Pondeville et al. 2014), we serendipitously created a transgenic line in which ectopic expression of the normally eye specific fluorescent marker gene was observed in larval, pupal and adult oenocytes. By cassette exchange of this marker with promoterless Gal4 activators, we tested whether the putative oenocyte enhancer could be trapped to drive Gal4 expression. Following successful enhancer trapping, we then examined the *in vivo* role of the two *An. gambiae cyp4g* genes by stable RNAi using UAS:hairpin DNA constructs regulated by the oenocyte Gal4 driver.

## Materials and Methods

### Plasmid construction

#### pBac-CFP

The docking lines were created via transformation of *Anopheles gambiae* embryos using a *piggyBac* vector containing the leucine-rich repeat protein immunity gene, LRIM1(Povelones et al. 2009), under the control of the *vitellogenin promoter*, *Vg*, with an eCFP 3xP3 eye marker gene (Horn et al. 2000), flanked by inverted *attP* sequences. The *vitellogenin* promoter, *Vg*, (AGAP013109) was removed from pBac[3xP3DsRed-AgVgT2-GFP] (Chen et al. 2008) with AscI and subcloned into pSLfa1180fa (Horn and Wimmer 2000) to give pslVg-GFP-BGH, which was subsequently digested with EcoRI and BamHI and the *Vg* promoter cloned into pSLfa1180fa.

BGH was amplified by PCR from pslVg-GFP-BGH by PCR with BamHI and NotI tagged primers (BGH-F, BGH-R) and subcloned into pJET1.2 (Thermo Scientific). *attP* was amplified from pBCPB+ (Groth et al. 2000) with XhoI and NotI tagged primers (attP-F, attP-R) and subcloned into pJET1.2. pslVg was digested with BamHI and XhoI, and the BGH and *attP* fragments, removed from pJET1.2 by BamHI /NotI and XhoI/NotI digests respectively, were cloned into pslVg simultaneously. LRIM1 (AGAP006348) with Strep and His epitope tags was amplified by PCR from pIEx10-LRIM1 (Povelones et al. 2009) with BamHI tagged primers (LRIM1-F, LRIM1-R) and subcloned into pJET1.2 (Thermo Scientific) before cloning into pslVg-BGH-attP. *attP* was released from pBac[3xP3-eCFPaf]-attP (Nimmo et al. 2006) by digestion with PstI, and *attP* reinserted in the opposite orientation by PCR amplification of *attP* from pBCPB+ with PstI tagged primers (attPPSTI-F, attPPSTI-R) to make pBac[3xP3-eCFPaf]-attPii. pslVg-LRIM1-BGH-attP was digested with AscI and cloned into pBac[3xP3-eCFPaf]-attPii to give pBac-CFP (Fig S1). All primers used are shown in Table S1.

#### pSL-attB-Hsp-Gal4(3xP3-dsRed)

RCME was carried out via PhiC31 recombination in the *attP* docking lines using a plasmid containing a minimal promoter upstream of a Gal4 coding region, and a dsRed 3XP3 marker (Horn et al. 2002) and flanked by inverted attB sites to allow cassette replacement via a double recombination event.

The minimal heat shock promoter Hsp70 was amplified by PCR from pUAST (Brand and Perrimon 1993) with NotI and EcoRI tagged primers (HSP-F, HSP-R), and subcloned into psl-LRIM-Gal4Δ and psl-LRIM-Gal4FF (Lynd and Lycett 2011) in place of the LRIM promoter. The 5’ attB site was amplified from pB*attB*[3×P3-dsRed2nls-SV40]*lox*66 using AflII and SpeI tagged (attB5-F, attB-R) primers and ligated into psl-Hsp-Gal4Δ and psl-Hsp-Gal4FF. A synthetic multiple cloning site was inserted into the 3’ unique BamHI site to provide EcoRI, SphI, NheI, NsiI and BglII sites. The 3’ attB site was subcloned into both Gal4 plasmids after PCR amplification from pB*attB*[3×P3-dsRed2nls-SV40]*lox*66 (Nimmo et al. 2006) using BglII and NsiI tagged primers (attB3-F, attB3-R). The 3xP3 promoter, dsRed gene and SV40 region were amplified from pBac[3×P3-dsRed] (Horn et al. 2002) using NheI and NsiI tagged primers (Red-F, Red-R) and subcloned downstream of the Hsp-gal4 cassette. Plasmids psl-attB-Hsp-Gal4[3xP3dsRed] and psl-attB-Hsp-Gal4GFY[3xP3DsRed] were constructed by subcloning Gal4 and Gal4GFY fragments from pLRIM-Gal4 and pLRIM-Gal4-GFY respectively (Lynd and Lycett 2011), into Psl-attB-Hsp-Gal4Δ[3xP3dsRed] in place of Gal4Δ using enzymes EcoRV and PspXI. All regions amplified by PCR were sequenced and all plasmids verified by diagnostic restriction digest (primers shown in Table S1).

#### pSL-attB-UAS-cyp16RNAi and pSL-attB-UAS-cyp17RNAi

To create a suitable UAS vector for expressing hairpin RNAi cassettes, the luciferase gene was removed from pUAS14-LUC (Lynd and Lycett 2011) by digestion with NcoI and XbaI and the plasmid was re-ligated using an oligo linker. The gypsy insulator sequence from pH-Stinger (Drosophila Genomic Resource Centre) (Barolo et al. 2000) was amplified by PCR with HindIII and SphI tagged primers (gyp1F, gyp1R) and inserted 5′ to the UAS. A second gypsy insulator sequence was inserted 3’ via PCR amplification using BamHI tagged primers (gyp2F, gyp2R). A multiple cloning site was then inserted into resultant plasmid by linker insertion (RINcoF, RINcoR) following EcoRI and NcoI digestion. Fusion PCR was carried out to create an *eYFP* marker under the control of the 3xP3 promoter with an SV40 terminator from plasmids eYFP-mem (Clontech), pBac[3×P3-dsRed]CP-Gal4-GFY-SV40 (Lynd and Lycett 2012) and pSL-attB-Hsp-Gal4[3xP3-dsRed] respectively. This cassette was cloned into the above UAS-gypsy intermediate plasmid with NsiI to give pSL-UAS14-gyp[3×P3-eYFP]. pSL-attB-Hsp-Gal4[3xP3-dsRed] was digested with NsiI and NotI and religated with an oligo linker (Link2F, Link2R) to give pSL-attB. pSL-UAS14-gyp[3×P3-eYFP] was digested with NotI and SpeI and the entire cassette ligated into pSL-attB to give pSL-attB-UAS14-gyp[3×P3-eYFP] (primers shown in Table S1).

To create the *cyp4g16* knockdown construct, genomic DNA (gDNA) was extracted from adult *Anopheles gambiae* females (G3 strain) using the method described in Livak (1984). A hairpin RNAi construct was then generated by direct amplification via asymmetric PCR of gDNA (Xiao et al. 2006) using exon 2 of *cyp4g16* (AGAP001076) to form the double stranded stem (198bp) and intron 2 to form the loop (579bp). PCR was carried out in a 25 μl reaction volume containing a final concentration of 1× HF buffer (ThermoScientific), 2.5 mM dNTPs, 0.4 μM forward primer and 0.2 μM bridge primer (primers shown in Table S1), 0.25 μl Phire DNA polymerase (ThermoScientific), and 1/200^th^ of a mosquito gDNA. Reaction conditions were 98°C for 30 sec, followed by 35 cycles of 98°C for 5 sec, 58°C for 20 sec, 72°C for 90 sec; followed by a final extension at 72°C for 10 min. The forward primer contained a 5’ NheI tag allowing the hairpin RNAi construct to be cloned into pSL-attB-UAS14-gyp[3×P3-eYFP].

To synthesize the *cyp4g17* knockdown construct, fusion PCR was carried out to fuse a 517bp region of exon 1 of *cyp4g17* (AGAP000877) and the proceeding 128bp intron (amplified from gDNA), to the reverse complement of the partial exon 1 fragment (amplified from cDNA). RNA was extracted from adult G3 females according to Tri Reagent manufacturer’s protocol (Sigma) and treated with Turbo DNase (Ambion). First strand cDNA was prepared from approximately 20 ng total RNA using Superscript III First-strand Synthesis System for RT-PCR (Invitrogen). Products from both amplifications were used as template for the final fusion PCR which was then cloned into pSL-attB-UAS14-gyp[3×P3-eYFP] with NheI and NcoI (all primer details in Table S1).

### Transformation of Anopheles gambiae

All *Anopheles gambiae s.s.* mosquitoes were reared under standard conditions and transgenic *attP* RCME docking lines were created by *piggyBac* mediated transformation of the G3 strain with the pBac-CFP using standard procedures (Lynd and Lycett 2012; Lombardo et al. 2009).

### Creation of Gal4 driver lines

Early embryos from transgenic *attP* docking line A14 were injected with buffer containing 350 ng/μl of one of the Gal4Δ, Gal4-FF, Gal4-GFY and native Gal4 protein variants of the attB-Hsp Gal4 driver plasmid and 150 ng/μl of intergrase helper plasmid, PKC40, as previously described (Lombardo et al. 2009; Pondeville et al. 2014). Surviving larvae were reared to adulthood and crossed to the G3 strain. G1 progeny were screened for fluorescent eye marker using a Leica MFLZIII microscope fitted with dsRed, YFP and CFP filter sets. Transgenic G1 larvae were pooled according to sex and crossed to the G3 strain. Isofemale lines were obtained from individual female lays and the G2 progeny interbred. DNA extractions and orientation PCR (see below) were carried out on the transgenic G1 parents and lines with a unique transgene cassette arrangement were maintained. A11 Gal4 lines were also generated in similar fashion to test the RCME efficiency using the native Gal4 attB-Hsp Gal4 plasmid.

### Creation of UAS p450 RNAi responder lines

Transgenic UAS responder lines containing the *cyp4g16* and *cyp4g17* RNAi cassettes were made by injection of the A11 embryos with pSL-attB-UAS-cyp16RNAi or pSL-attB-UAS-cyp17RNAi plasmid (350ng/ul) together with the intergrase helper plasmid, PKC40 (150ng/ul) as described previously. G1 larvae positive for eYFP, indicative of an integration event, were used to generate isofemale lines and screened by PCR to determine the orientation of the cassette, as described below.

### Determination of insertion arrangement

A series of four diagnostic PCRs were carried out to determine both the orientation of the *attB* cassettes and, for mosquitoes having both markers, the relative position to the *attP*-*CFP* cassette, using one external primer and one internal primer (see Fig S3A to describe potential orientations and Table S1 for primers used). DNA from a single G1 adult was extracted using a Qiagen DNeasy Blood & Tissue Kit as per manufacturer’s protocol, and DNA eluted in 200μl of ddH20. PCR was carried out using the manufacturer’s recommendations for Phire Hot Start II Polymerase (NEB) using 1 μl of DNA and a final primer concentration of 100nM each, with an annealing temperature of 58°C for 20 seconds, and an extension time of 90 seconds. PCR products were sequenced directly to confirm a precise recombination event at the *attP*/*attB* junction.

### Southern Blots

DNA was extracted from at least 30 male and female adult mosquitoes using Qiagen GenomicTip 20/G columns as described in manufacturer’s standard protocol. 9 μg of genomic DNA was then digested to completion with EcoRI (NEB), purified on diatomaceous earth (Carter and Milton 1993) (Sigma) and run on a 0.8% agarose gel. DNA was transferred to a nylon membrane (Lycett et al. 2004). Two probes were amplified from a plasmid template targeting the pBac arms (see Table S1 for primers) and labelled with CTP α-32P 3000Ci/mmol (Perkin Elmer) using Klenow Enzyme (NEB) and random nonamers (Sigma). Hybridization, washing and exposure to X Ray film were performed as described previously (Lycett et al. 2004).

### Whole-mount larval abdomen immunostaining

Abdominal integuments were dissected from 4^th^instar larvae and fixed and stained as described previously (Lycett et al. 2006). The rabbit affinity purified CYP4G16 and CYP4G17 antibodies (Ingham et al. 2014; Balabanidou et al. 2016) were used as a 1/500 dilution and detected with goat anti-rabbit antibody (Alexa-Fluor 488; Molecular Probes; 1/1,000). Nuclei were stained with ToPRO 3-Iodide (Molecular Probes). Images were obtained on a Leica SP8 confocal microscope.

### Analysis of enhancer trapped gal4 expression

To investigate enhancer driven gal4 expression, crosses were made between the HSP-Gal4 mosquito lines (marked by dsRed) and the UAS mosquito line, Wnd, containing *luciferase* and e*YFPnls* reporter genes (marked by eCFP) (Lynd and Lycett 2012). At least 10 females were used per cross, with at least double the number of males. Larval progeny were screened for expected inheritance of both red and cyan fluorescent eye marker proteins. Adults were examined at three and seven days after emergence. Dissections were carried out using a Leica MFLZ III microscope using YFP, dsRed and CFP filters. Images were taken using an Olympus BX60 microscope fitted with a Nikon DSU2 camera, or through a dissecting microscope fitted with a Nikon P5100 digital camera.

### Luciferase assays

Adult mosquitoes were anesthetized with CO2 when three to five days old and dissections carried out in PBS. Whole and dissected mosquito larvae and adults were transferred to 200μl of luciferase passive lysis buffer (Promega). Samples were homogenized, and the supernatant used immediately for luciferase assays using a Promega Luciferase Assay Kit (E1500) and a Lumat LB 9507 tube luminometer. At least six replicates for each assay were carried out.

### Investigation of knock down phenotype

Crosses were made between the oenocyte specific Gal4 driver line (DsRed), A14Gal4, and the UAS *cyp4g16* and *cyp4g17* RNAi mosquito lines (YFP) originating from lines carrying integrated hairpin constructs into the *attP* docking site of line A11. Larvae and adults of the resulting progeny carrying both markers, designated 16i and 17i henceforth, were examined for various phenotypes using siblings arising from the same cross without the UAS cassette (dsRed only) as negative controls.

### qRT-PCR of cyp4g16 and cyp4g17

RNA was extracted from different mosquito stages and treated with DNase as above. First strand cDNA was prepared from total RNA using Superscript III First-strand Synthesis System for RT-PCR (Invitrogen). Transcript abundance of the *cyp4g16* and *cyp4g17* genes were examined using two sets of primers for each gene in order to monitor reliability of amplification (details in Table S1). One primer of each pair was designed to span an exon-exon junction. The PCR efficiency, dynamic range and specificity of the primer pair were calculated from running a standard curve over a five-fold dilution series of cDNA. Transcript levels were normalized against amplification of the *An. gambiae* ribosomal protein genes, S7 (AGAP010592) and Ubiquitin (AGAP007927) (Jones et al. 2013). PCR was carried out in a 20 μl reaction volume containing 10 μl Brilliant III SYBR Green Master Mix (Agilent), 500 nM of primers, 1μl cDNA (diluted 100-fold). qPCR was performed on the MXPro qPCR system (Agilent) and reaction conditions were 95°C for 10 min followed by 40 cycles of 95°C for 10s and 60°C for 10s. Melt curves were performed after each PCR to ensure the specificity of the qPCR. Three technical replicates were run for each sample and a minimum of four biological replicates carried out.

### Western Blots

Early pupae (less than 2hours old) were extracted in 1x Laemlli buffer (BioRad) with 100mM DTT. Proteins were separated on 10% SDS-PAGE with a Tris-glycine running buffer (192 mM glycine 25 mM Tris, 0.1% SDS) and transferred to polyvinylidene fluoride (PVDF) membrane. Filters were blocked with 3% powdered milk in PBST (PBS+0.1% Tween-20) for 1h at room temperature and then incubated with anti-CYP4G16 (1:200 dilution (Balabanidou et al. 2016)), anti-CYP4G17 (1:200 (Ingham et al. 2014)) or anti-alpha tubulin (1:500 Sigma) antibodies in 3% milk-PBST for 1 h at room temperature. Membranes were washed four times in PBST for 5 min, then incubated with anti-rabbit (1:40,000) or anti-mouse (1:10,000) secondary antibodies conjugated to peroxidase. Filters were developed with the Western blot Chemiluminescence Reagent Plus Kit (Renaissance) and exposed to X-ray films.

### Gas chromatography-mass spectrometry analysis

Cuticular hydrocarbons of single 16i and 17i one day old female adults or female pupae collected within 5 hours of larval/pupal transition, together with identically reared sibling A14Gal4:+ mosquitoes, were extracted by a 5 minute immersion, with gentle agitation at room temp, in 50 μl hexane (Sigma-Aldrich) containing 10 ng/μl octadecane (Sigma-Aldrich) internal standard. Hydrocarbon identification and quantification was performed by gas chromatography-mass spectrometry (GC-MS). The mass spectrometer employed was a Waters GCT and the GC column was a 30 cm long, 0.25 mm internal diameter, 0.25 μm film thickness BPX5 (SGE). The oven temperature gradient was 70°C to 370°C at 10°C/minute and the carrier gas was helium (BOC) at a flow rate of 1 ml/minute. The injection volume was 1 μl. The MS scan range was *m/z* 40 to 450 Da in scan time 0.6 s. Peak identification was by a combination of retention time, library searches using the NIST mass spectrum library supplied with the instrument and reference to published spectra. Peak areas were measured manually using the peak integration tool in the Waters Mass Lynx software. A representative annotated trace is provided in File S1. The total amount of hydrocarbon present was calculated by summing all of the peak areas detected relative to the internal standard. Statistical analysis of total CHC was performed by ANOVA with Tukey HSD correction. The relative quantity of individual hydrocarbons present in each mosquito sample were compared in two ways. 1) relative to the internal standard, by dividing the area of each extracted hydrocarbon peak to that of the internal standard to give a quantitative value to each hydrocarbon and 2) relative to the total hydrocarbon content in each mosquito by dividing each area by the sum of all areas to give a fraction of the total (statistical analysis provided in File S2 and File S3 for normalization against Internal Standard and total CHC respectively).

### Desiccation Assays

10-13 mosquitoes aged two to five days were placed in 70 ml transparent polystyrene pots and a netting mesh lid attached with an elastic band. Pots were then placed in 300 ml transparent polystyrene desiccation chambers with 10g silica gel desiccant, sealed with a screw on cap and placed in a 32°C incubator. Desiccant is thought to reduce humidity to < 10%, (Gray and Bradley 2005). 7 replicates for 16i and A14A1 sibling females, and 6 replicates for 16i and A14A1 sibling males were performed. Survival was measured by the ability to stand, cling to walls or fly and was scored at the indicated times. Statistical analysis was performed by Cox regression.

### Data and Reagent Policy

All extant mosquito lines and novel plasmids described in the manuscript are available for distribution. Supplementary data deposited at Figshare includes Schematic of the constructs used for transformation Figure S1, Summary of generation of docking lines Figure S2, Phenotypic characterisation of RCME lines Figure S3, Dissection of Female Adult and Pupa derived from A14Gal4 x UAS-mcherry cross Figure S4 Images of A11Gal4 X UAS-nlsYFP progeny Figure S5 CYPG16 and CYP4G17 larval expression Figure S6, Schematic of the constructs used for gene knockout Figure S7, Western analysis of 16i,17i and A14A1 early pupae, Figure S8, Plots of relative abundance of CHCs with respect to internal standard Figure S9, Plots of relative abundance of CHCs with respect to total CHC Figure S10. Sequences of primers described in Materials and Methods Table S1, Summary table of RCME experiments performed with different Gal4 constructs in A14 docking line Table S2, Representative, annotated GCMS CHC trace from individual mosquitoes File S1, Statistical analysis of CHC normalized to Internal standard File S2, Statistical analysis of CHC normalized to Total CHC content File S3.

## Results

### Generation of oenocyte driver line

#### Oenocyte expression in docking line A14

We created a series of *piggyBac* based PhiC31 docking strains of *An. gambiae*, carrying inverted *attP* sites flanking the cyan fluorescent protein (*eCFP*) selectable marker and the *LRIM*1 (Povelones et al. 2009) gene controlled by the blood meal inducible Vitellogenin promoter (Chen et al. 2008)(Fig S1A). From 10 isofemale lines produced (Fig S2A), four carried insertions into single genomic sites, as indicated by inverse PCR (Fig S2B). All lines had the expected expression of eCFP in eyes and neuronal tissues, and six out of eight examined expressed the tagged *LRIM1* transcript (Fig S2C) 48hrs after bloodfeeding. Western analysis indicated that three insertion lines with single copy insertions, A11, A14 and BB57, expressed the tagged LRIM1 protein after bloodfeeding (Fig S2D).

However in one line, A14, distinct eCFP fluorescence was also readily detected in oenocytes in larval and pupal stages (Fig 1), suggesting that one or more local genomic enhancers were directing eCFP expression to these abdominal cells. Sequencing around the transposon insertion site in A14, indicated that integration had occurred at an intergenic site on the 3R chromosome (Fig S2E), greater than 30Kb away from the nearest annotated gene in the PEST genome (Vectorbase P4.12). Since we created the A14 line to carry inverted *attP* sites surrounding the marker and effector cassette, we surmised that recombinase-mediated cassette exchange (RCME) with a promoterless Gal4 construct would bring the transctivator under control of the enhancer to produce oenocyte specific driver lines.

**Figure 1.**
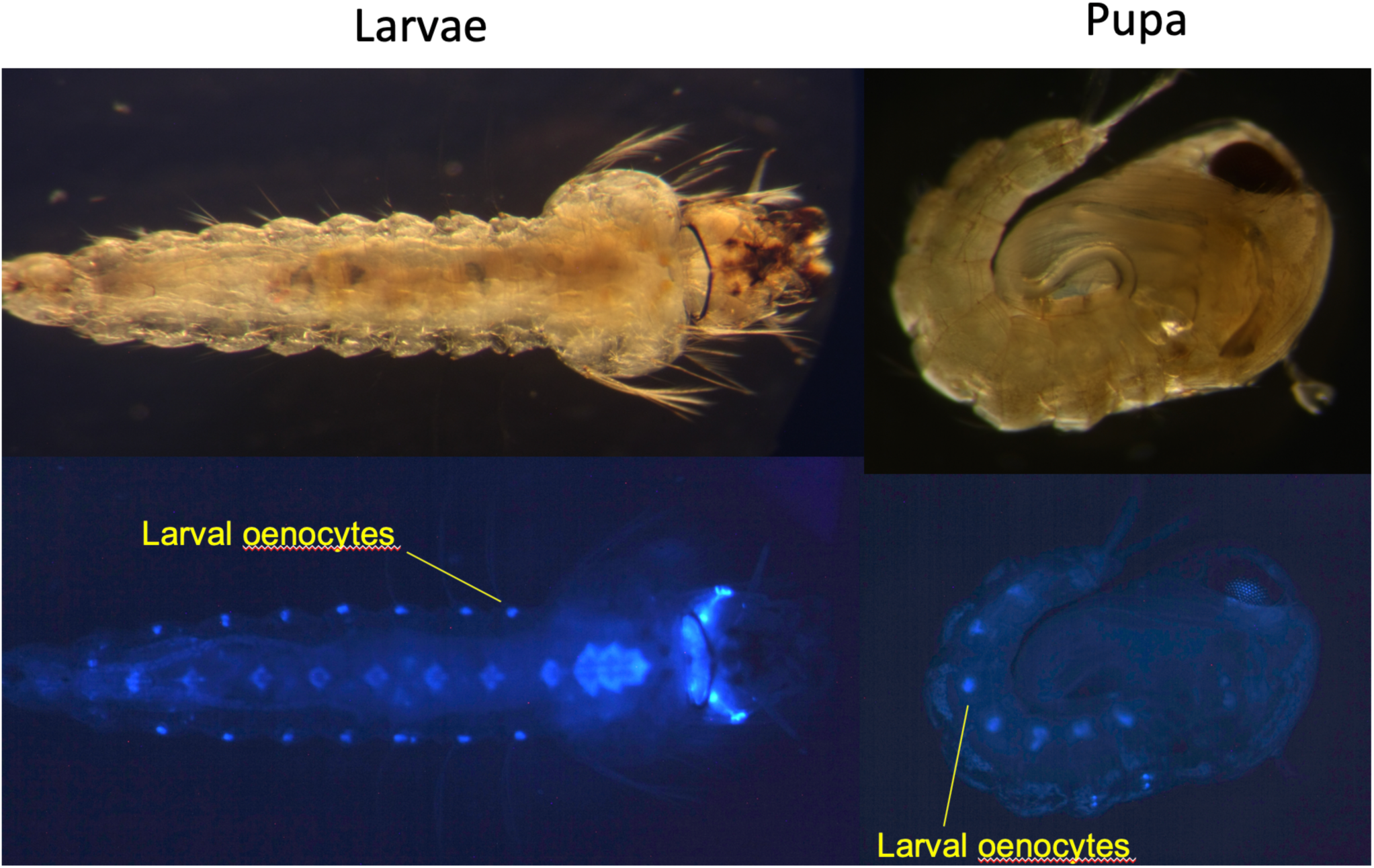
Oenocyte-specific expression of ECFP marker in A14 larvae and pupa Oenocyte-specific expression of eCFP in larva and pupa in docking line A14 presumably caused by a local oenocyte enhancer near to pBac-CFP [attP-LRIM1] integration site. Upper figures are bright field images and lower figures are images taken through CFP filter on stereofluorescent microscope. LO indicates larval oencoyctes which persist during young pupae. ECFP expression in eyes and ventral nerve cord are characteristic of 3xP3 expression.

### RCME is efficient in mosquitoes

To initially develop an efficient oenocyte driver, we examined the efficacy of four alternative Gal4 constructs marked with dsRed, by targeting cassette exchange into A14 embryos. Previous cell culture analysis indicated that these alternative Gal4 constructs, native Gal4, Gal4-delta, Gal4-GFY and Gal4-FF generate different transcriptional activation potentials (Lynd and Lycett 2011), and so may produce a graded range of oenocyte-specific expression levels. As indicated (Table S2), between 6 and 155 dsred positive larval progeny were successfully generated for each construct. However, viable adults were only obtained using the native Gal4 (>100 adults) and Gal4-FF (1 adult from 6 dsred positive F1 larvae) transactivators.

Should canonical cassette exchange of the dsred marked Gal4 driver have occurred in the eCFP marked A14 line, the red marker would replace the blue marker, in two potential orientations. Whereas integration into a single *attP* site, in four possible orientations, would conserve the *eCFP* marker in this locus, and thus F1 progeny would be doubly marked (this is illustrated in Fig S3A). In total seven isofemale native Gal4 lines were bred from surviving F1 progeny; three exchange and four integration lines. PCR analysis indicated that five of the six possible integration patterns were detected within these lines (Table S2). Southern hybridization on the representative A14Gal4 line (A1 line kept for further characterisation) confirmed cassette exchange into the expected genomic site and orientation (Fig 2A and B).

**Figure 2.**
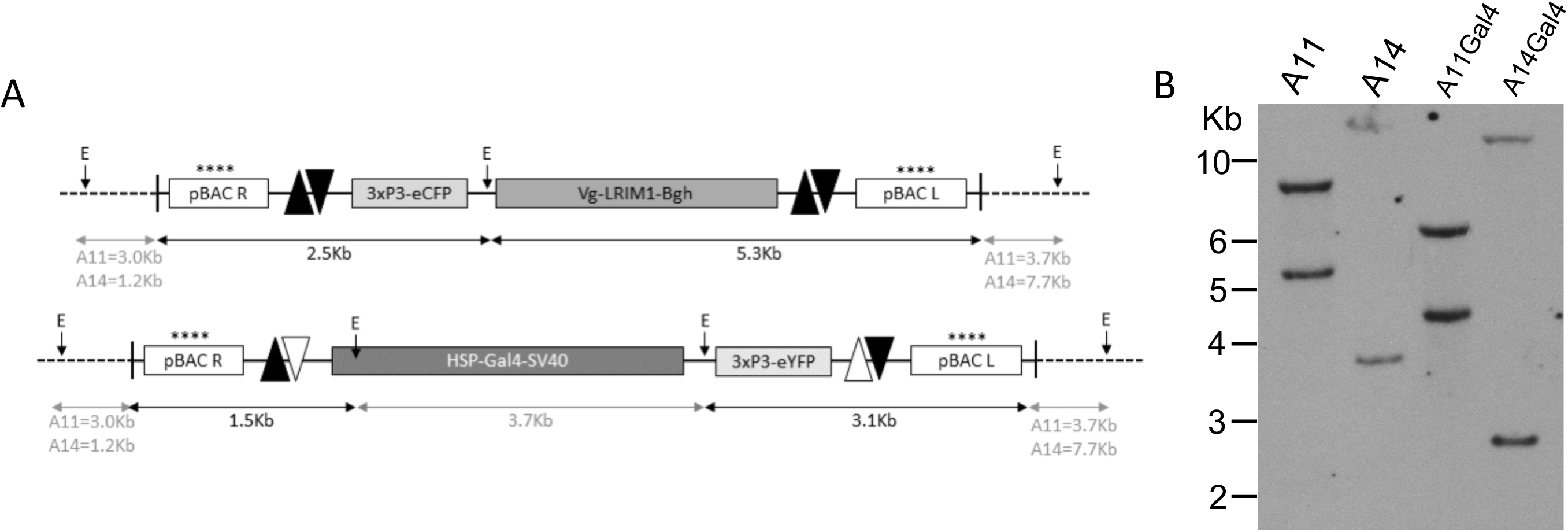
Schematic of *piggyBac* and RCME genomic integrations and Southern analysis A: schematic of PBac-CFP [attP-LRIM1] integrated by RMCE into genomic DNA of A14 and A11 lines and orientation of exchanged attB HSP-Gal4 cassettes. ****, pBAC L and pBAC R probes; E, EcoRI restriction site; Paired black triangles, *attP* sites; mixed black and white paired triangles, attL and attR sites resulting from phiC31-mediated recombination. Distance from pBac integration sites to flanking genomic EcoRI sites were estimated from the southern blot as A11 3.0Kb and 3.7Kb, A14 1.2Kb and 7.7Kb. B: Southern blot showing single copy integration (two probes – two bands) and shift in expected size of hybridizing fragments following exchange of attB HSP-Gal4 into A11 and A14 genomic sites.

Since RCME efficiency had not previously been assayed in mosquitoes, we also tested a second control docking line, A11, that displayed the typical expression of the 3xP3 marker. Into the A11 line we exchanged the same native Gal4 construct and obtained 21 isofemale lines from 638 embryos injected. Diagnostic PCR was again used to define orientation of insertion in six of these lines (Table S2) and Southern blotting of an A orientation isofemale line confirmed that RCME had occurred at the expected site due to the expected change in restriction enzyme pattern (Fig 2A and B) compared the parental line.

### Oenocyte enhancer is trapped by RCME

In the A14 native Gal4 trap lines, weak red fluorescence was detected in oenocytes following exchange in the A14A1 and A14A2 lines (Fig S3B). To examine whether Gal4 expression was also under oenocyte enhancer control, the A14 Gal trap lines with alternative orientations of insertion (including alternative isofemale lines with the same molecular orientation, A14A1 and A14A2) were crossed to a 3xP3 *CFP* marked responder line carrying nuclear localised yellow fluorescent protein (nlsYFP) and luciferase genes both under control of the upstream activation sequence (UAS), to allow qualitative and quantitative analysis, respectively (Lynd and Lycett 2012). In progeny of these crosses, nlsYFP was visible in larval (Fig 3B), pupal and adult oenocytes (Fig 3D), indicating that enhancer trapping was achieved. Furthermore, through Gal4 enhanced expression we could trace the development of adult oenocytes through 4^th^ instar larval (Fig 3F) and pupal stages as they contained nlsYFP. The same oenocyte expression profile was obtained in progeny following A14Gal4A crossing with a separate UAS-mCherry line (Adolfi et al. 2018) (Fig S4). Progeny from crosses between the A11Gal4A line with the UAS-nlsYFP line are shown in Fig S5 and lack expression in the oenocytes.

**Figure 3.**
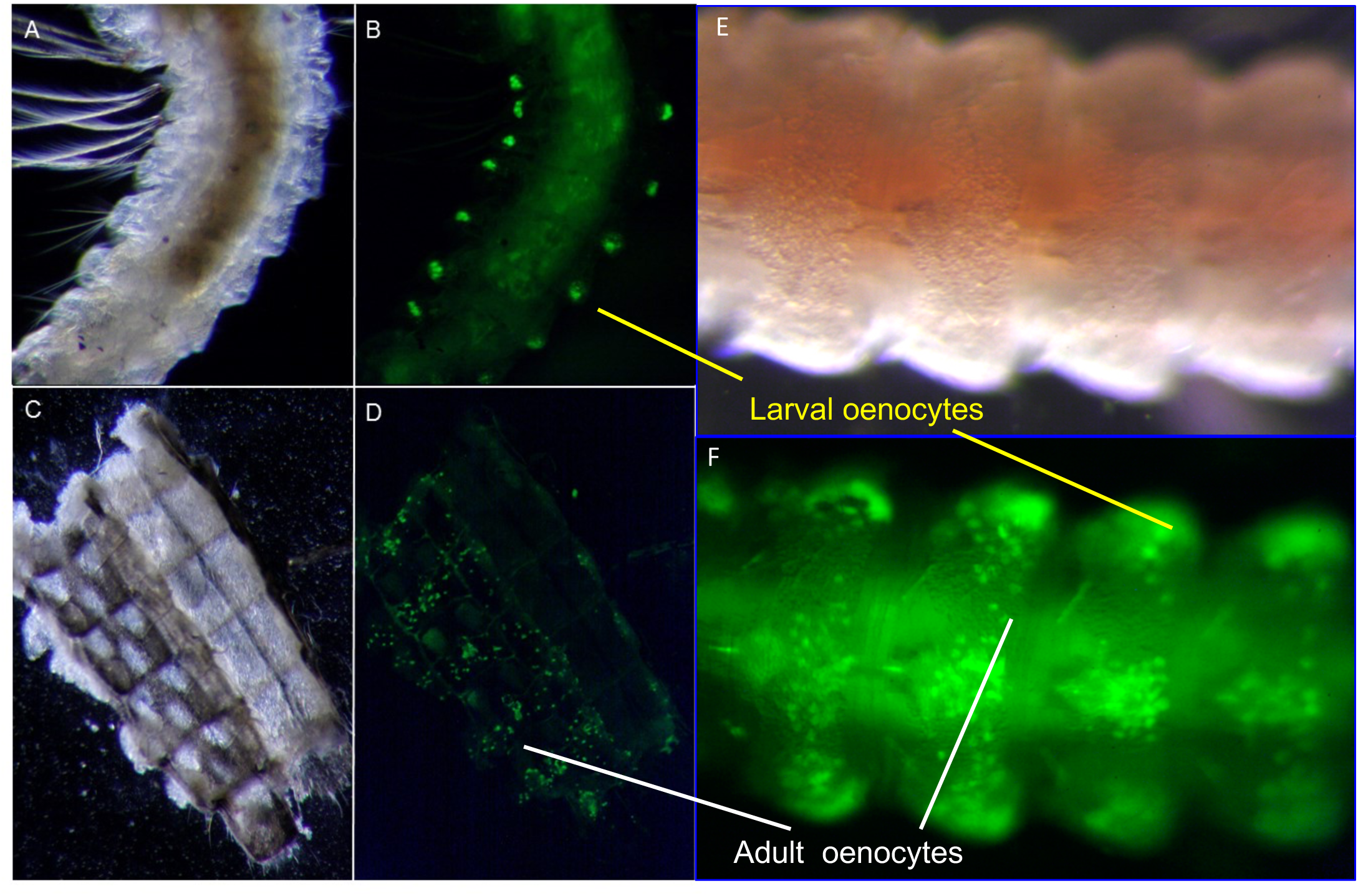
Oenocyte Enhancer trapping in larval and adult oenocytes Images of yellow nuclear florescence in progeny of crosses between A14Gal4 driver line and the UAS responder line containing *eYFPnls* reporter genes. A and B; show YFP expression in the oenocytes located in the abdomen of 3^rd^ instar larvae. C and D; show bright field and fluorescent images of the dissected adult abdomen integument. YFP expression is visible in the larval oenocytes (LO) and adult oenocytes (AO). E and F show bright field and fluorescent images of 4^th^ instar larvae showing the development of adult oenocytes on the ventral side of the integument.

Quantitative analysis of the different orientation A14Gal4 lines (Table S2) by luciferase activity analysis indicated that in both 4^th^ instar larvae and adults, the cassette orientation (Fig S3A) had a significant effect on reporter gene expression. Those with orientation A had three-fold more activity than those with orientation B at larval stages (Fig S3C), and nine-fold greater expression at adult stage (Fig S3D). Further analysis of the A14A1 line indicated that luciferase activity increases 100-fold during larval development, plateaus during metamorphosis, before dropping four-fold in adult stages (Fig S3E).

### Expression of 4Gs in immature stages

To examine the utility of the driver line for oenocyte-specific knockdown of CHC regulating enzymes, we targeted two genes, *cyp4g16* and *cyp4g17*, that previous work had established were highly expressed in adult oenocytes (Ingham et al. 2014; Balabanidou et al. 2016). To determine the expression profile of native mosquito *cyp4g*s in immature stages, we performed qPCR analysis on RNA extracts from larval and pupal samples collected at defined timepoints following moulting. *cyp4g16* was shown to be highly transcribed relative to housekeeping gene controls (delta C^T^: Fig 4A) throughout the 3^rd^ instar and significantly increased (up to eight-fold) during 4^th^ instar and pupal stages (Fig 4A). Whereas the *cyp4g17* transcript was at the minimal limits of detection in 3^rd^ instar stages, and expression massively increased (up to 10,000-fold) after 8 hours of the early 4^th^ instar stage, before decreasing during pupal moult, and again increasing significantly at later pupal stages (Fig 4B). Immunostaining confirmed that both CYP4G proteins are readily detected in 4^th^ instar larval oenocytes. (Fig S6 A,B).

**Figure 4.**
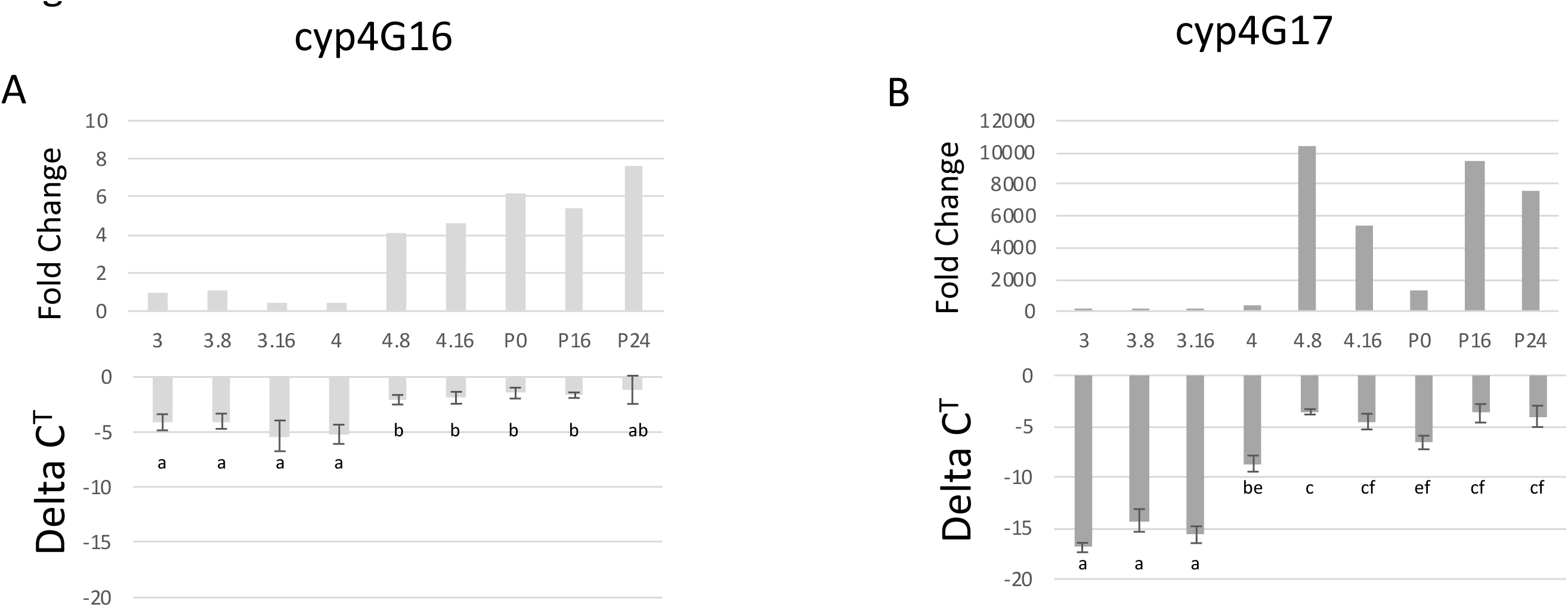
QRT-PCR analysis of *cyp4g16* and *cyp4g17* during larval and pupal stages Lower graphs show mean delta C^T^ differences in expression compared to the housekeeping controls to clearly demonstrate the higher constitutive expression of *cyp4g16* in earlier stages, with error bars indicating standard error of the mean. Upper graphs are transformed C^T^ data that shows the relative fold change in expression (delta delta C^T^) at each stage compared to newly moulted 3^rd^ instar larvae to demonstrate the large relative induction of *cyp4g17* expression at later stages. Assays were performed in triplicate on four independent samples. 3 refers to newly moulted 3^rd^ instar larvae, 3.8: 3^rd^ instar larvae aged 8 hours from moult, 3.16: 16 hours from moult, 4: newly moulted 4^rd^ instar larvae, 4.8 4^rd^ instar larvae aged 8 hours from moult, 4.16 16 hours from moult, P0: newly moulted pupa, P16: Pupa 16 hours from moult, P24: Pupa 24 hours from moult. Statistical analysis is performed on C^T^ data and those values sharing letters do not show significant difference between themselves at p<0.05 cutoff by Welch test. The fold change data is simply transformed C^T^ data and statistical analysis should be taken from the raw C^T^ data.

### Oenocyte Gal4-driven knockdown of *cyp4g16* or *cyp4g17* produces different survival phenotypes

UAS responder constructs were created carrying inverted repeats of DNA from each *Cyp4g* gene, separated by their natural intron sequences, as shown in Fig S7. These were exchanged by RCME into the control A11 *attP* line, and distinct eYFP isofemale responder lines carrying insertions of UAS-*Cyp4g16* and UAS-*Cyp4g17* hairpins in the same orientation were generated. No deleterious phenotype has been observed in these lines and they have been maintained in colony for 4 years since production.

However, following crossing to the A14 driver line, the resultant transheterozygote progeny undergo death at distinct pupal stages depending on the specific RNAi construct present. Progeny expressing *cyp4g16* RNAi in oenocytes (16i) show high mortality (above 90%) at early-mid pupal stages (Fig 5 A), which is reduced if grown at low density (<200 larvae in a 30×30×5cm tray - in a typical experiment we observed around 60% death at early pupal stage, Table S3). The progeny expressing *cyp4g17* RNAi in oenocytes (17i) develop till late pupal stage and have high mortality (>90%) in pharate adults or as the adults emerge from the pupal case (Fig 5B, Table S3). Altering growth conditions does not increase 17i survival. Homozygous A11 and sibling heterozygous A14Gal4 mosquitoes typically show greater than 90% survival during 4^th^ instar larval to adult transition (Table S3).

**Figure 5.**
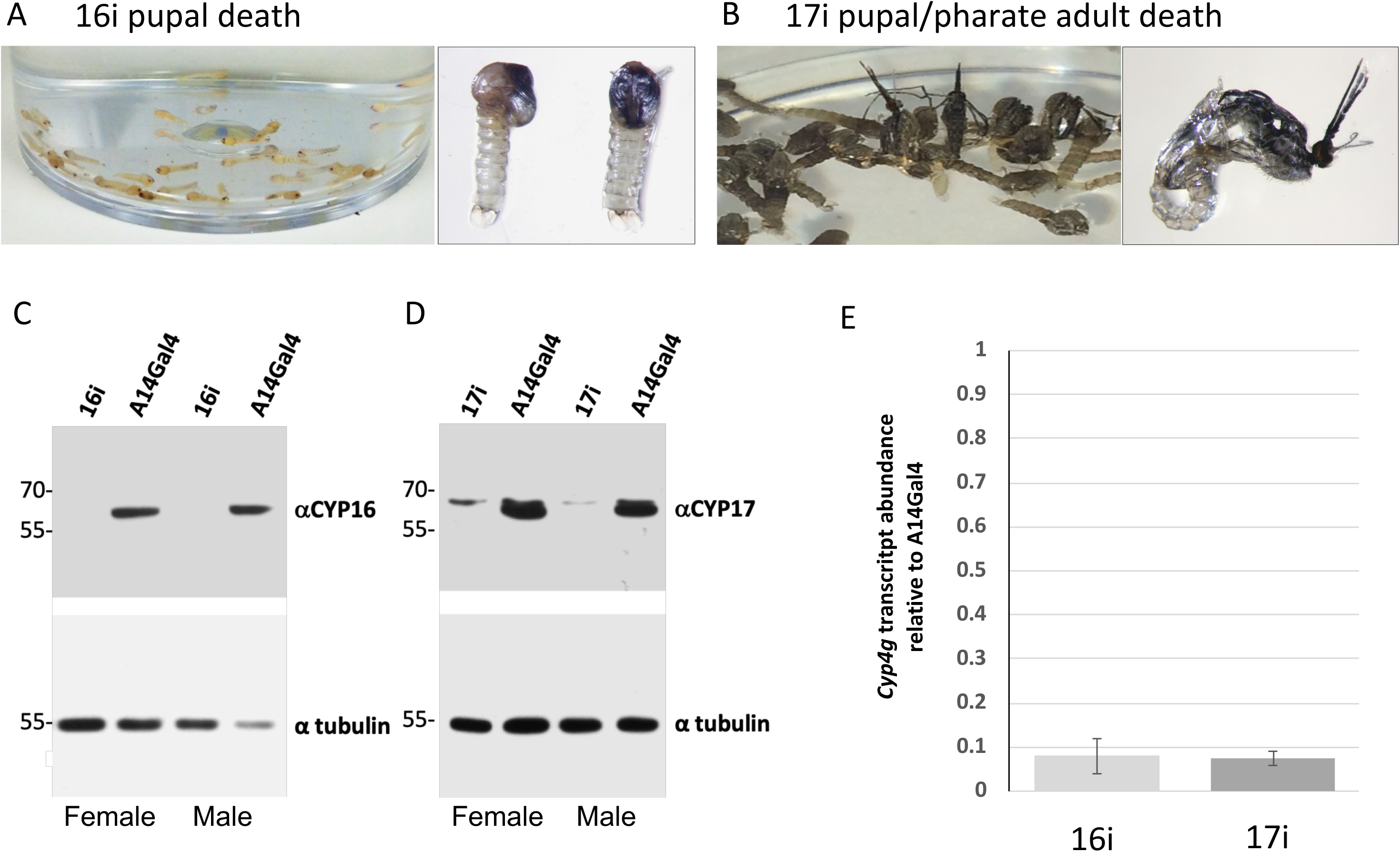
Analysis of 16i and 17i knockdown efficiency and lethality phenotype A and B Images displaying the different temporal pupal phenotype deaths occuring with 16i and 17i knockdown mosquitoes respectively. A: 16i pupae have a high mortality at early pupal stage characterised by sinking in water (right) and elongation of the pupae (left) at death rather than the characteristic comma shape. B: 17i death is characterized by very late pupal death (left) usually after the pupal case is cleaved at eclosion or during adult emergence (right). C and D: upper western analysis of knockdown of CYP4G16 in 16i mosquitoes and of CYP4G17 in 17i mosquitoes using the respective polyclonal antibodies. C and D: lower loading control of same filters probed with anti-alpha tubulin antibody. E: Analysis of transcript knockdown of the respective *cyp4g* by qRT-PCR analysis of *cyp4g16* and *cyp4g17* in 16i and 17i pupae compared with A14Gal4 (normalized to 1) sibling control mosquitoes that carried the oenocyte driver but lack the UAS-hairpin constructs.

### Stable RNAi efficiently knocks down expression of *cyp4g* genes

To examine efficiency of knock down of the *cyp4g* genes, western analysis and qPCR was performed on early pupae prior to death in 16i and 17i mosquitoes. Western analysis revealed an absence of CYP4G16 in 16i mosquitoes (Fig 5C). Similarly, the major protein corresponding to CYP4G17 was absent in 17i pupal extracts (Fig 5D). In concordance with lack of protein, transcript levels were reduced by 90% in both knockdowns relative to Gal4 driver line, A14 (Fig 5E). Western analysis also indicated that knock down was specific to the respective CYP4G proteins, since the level of CYP4G16 was similar to controls in 17i knockdown mosquitoes, and *vice versa* (Fig S8).

### CYP4G16 and CYP4G17 have a synergistic role in CHC synthesis in pre adult stages

Total CHC analysis of early pupae hexane cuticle extracts by gas chromatography– mass spectrometry (GC-MS) demonstrated that 16i or 17i mosquitoes both had a significant reduction of approximately 80% compared to controls (Fig 6A). Whereas, in surviving adults, this reduction in total CHC content was approximately 50% in both knockdowns (Fig 6B). However, there was no significant difference in total CHC between the 16i and 17i mosquitoes at either stage.

**Figure 6.**
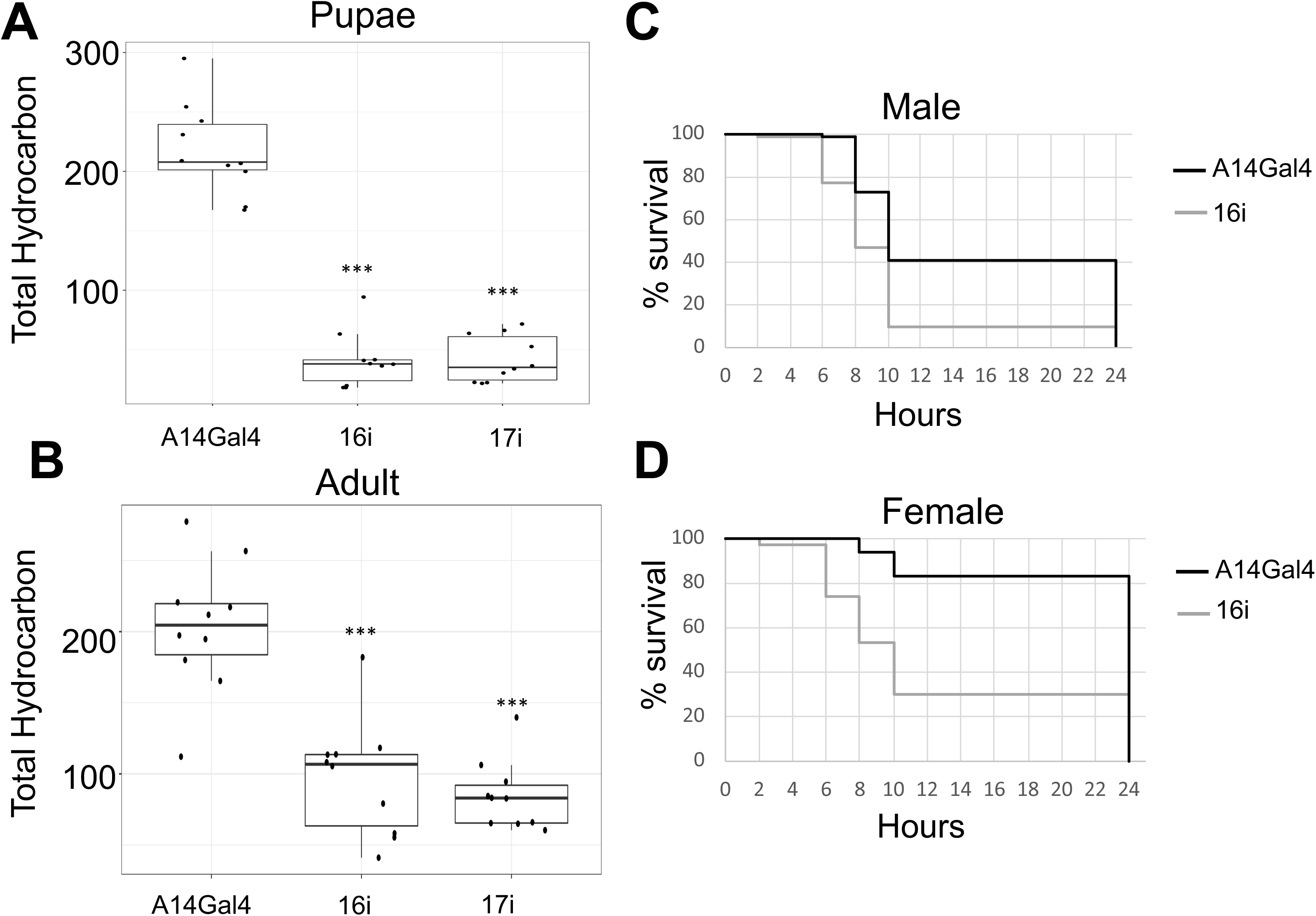
Total CHC content in female 16i and 17i mosquitoes and desiccation tolerance of 16i males and females. A: histogram illustrating total CHC content of female pupae collected within 5 hours of larval/pupal transition. B: histogram illustrating total CHC content of one day old female adults. Black bars indicate the median total CHC content per mosquito, black circles are values per mosquito. Boxes are 25^th^ and 75^th^ percentiles and whiskers + and – 1.5 x interquartile range. *** P<0.001 calculated by ANOVA followed by Tukey’s HSD (Honestly Significant Difference) for pairwise comparisons. C and D: survival curves for 16i and A14Gal4 male (C) and female (D) mosquitoes exposed to low humidity. Statistical analysis performed by Cox regression (Analysis and P values indicated in results text).

Separating the analysis down to the relative peak areas of individual hydrocarbons (with respect to the internal standard), no CHC species showed a significant difference (File S2) in abundance between the 16i and 17i knockdowns at the early pupal stage (Fig S9). In contrast, all individual CHCs (except nonadecane) were significantly reduced (p<0.05) in 17i pupae compared with controls (Fig S9). Most CHCs were also significantly reduced in 16i pupae compared to controls, although nonadecane p=0.55, icosane p=0.08, triacontane p=0.56 and hentriacontane p=0.08 were exceptions. At the adult stage, nonadecane was significantly reduced in 16i mosquitoes compared to 17i, whereas methyl-hentriacontane was significantly reduced in 17i compared to 16i mosquitoes. The major adult peaks, penta-, hepta- and nona-cosane were significantly reduced in both 16i and 17i mosquitoes compared to controls, as were hexacosane and hentriacontane. Additional significant differences between 17i and control mosquitoes were observed with octacosane, and methyl-nonacontane and methyl-hentriacontane (Fig S9).

Analysis of the relative proportion that individual CHCs comprise of total CHC content indicate that in control pupa, four odd-number chain alkanes provide over 70% of all hydrocarbons (henicosane 22%, tricosane 15%, pentacosane 17.5%, heptacosane 18%) (Fig S10). The relative proportions of all four major CHCs are significantly reduced (p<0.05) in both 16i and 17i pupae compared to controls (File S3). In addition, whilst the relative proportion of the two methyl branched chain CHCs (methyl-nonacosane and methyl-hentriacontane) are also reduced in 16i pupae (p<0.01, p<0.001), only the latter (p<0.01) is also reduced in 17i pupae compared to controls. Contrastingly, nearly all of the other minor peaks show significant relative increase in the 16i and 17i knockdown pupae compared to controls. When directly comparing the two knockdown mosquitoes, the relative proportion of hepta- and nona-cosane and the two methyl branched chains are significantly lower in 16i, whereas octacosane and tricontane are significantly lower in 17i pupae than 16i.

In control adults, there is a shift to higher molecular weight CHCs compared to control pupa with two odd chain alkanes providing over 50% of all hydrocarbons (heptacosane 35%, nonacosane 19%). In 16i knockdowns, both of these CHCs are significantly reduced in relative proportion compared to control. Whereas heptacosane and methyl-hentriacontane are significantly lower in 17i mosquitoes compared to control. Again the majority of the minor peaks show increases in relative abundance in both knockdowns. Comparison of the 16i and 17i knockdowns indicated that nonadecane and nonacosane were significantly reduced in 16i knockdowns, whereas methyl-hentriacontane (p<0.001) was significantly reduced in 17i knockdown. There was also a non significant trend (p=0.06) in relative reduction in methyl-nonacosane in the 17i knockdown.

### *cyp4g16* knockdown increases susceptibility to desiccation

Due to the high mortality in 17i pharate adults, the effect of CYP4G depletion on desiccation tolerance was only assessed in 16i adults. The time course of survival of 16i males and 16i females versus co-reared, age matched, sibling Gal4 controls when exposed to a low humidity environment is illustrated in Fig 5C and Fig 5D, respectively. Cox proportional hazards analysis of the time to mortality data was initially fit to a model comprised of three terms, for sex, genotype, and sex by genotype interaction effects, respectively. The interaction term was found to be non-significant so was removed from the model. This analysis suggested highly significant effects of both sex (p < 0.001) and genotype (p < 0.001) on the risk of death. In other words, males show significantly different survival to desiccation compared to females, and gene knockdown has a significant effect on survival time irrespective of sex. Moreover, analysis of individual sexes, males and females indicated that both 16i males (p < 0.001) and 16i females (p < 0.001) were significantly more sensitive to desiccation than their respective same sex controls.

## Discussion

Following serendipitous detection of oenocyte-specific fluorescence in one of a panel of RCME docking lines, we have successfully performed Gal4 based enhancer trapping in *An.gambiae* for the first time. Previous work has shown that enhancer trapping was feasible in mosquitoes through mobilisation of tagged transposable elements, and the subsequent changes in tissue specific expression of reporter genes (O’Brochta et al. 2011). Further work by this group developed enhancer trapping (O’Brochta et al. 2012) and more recently large scale promoter/gene traps in *An. stephensi* (Reid et al. 2018). We have taken a modified approach through the use of RMCE to insert alternative Gal4 genes nearby a putative enhancer. Although there is, as yet, no annotation for a gene closer than 30kb from the A14 insertion site, oenocyte-specific regulation is achieved by cassette exchange demonstrating that the locus contains DNA regions which interact with the minimal promoter present on the Gal4 driver. The work has thus confirmed that RCME is efficient in *An. gambiae* (Hammond et al. 2016). We have also generated a docking line, A11, into which RCME and subsequent Gal4 dependent gene silencing phenotypes were demonstrated with UAS-regulated hairpin RNAi constructs.

To develop the oenocyte driver, we assessed four variants of Gal4, each carrying alternative transactivators. These were shown previously to have graded activation potentials with Gal4Δ> GFY > Gal4 > FF in transfected *An. gambiae* cells (Lynd and Lycett 2011). In these cells, the relative activation potential varied 20-fold between the four transactivators, with Gal4Δ being 10-fold more active than Gal4. RCME derived dsred positive G1 larvae were obtained with each construct, but only those generated with native Gal4 transactivator surviving in large numbers to the adult stage, with the majority of those derived from Gal4Δ and GFY dying at early larval instar. This is in contrast to previous work with GFY combined with a gut specific carboxypeptidase promoter which gave very high transformation rates and robust homozygous adults that are still in colony (Lynd and Lycett 2012). This would suggest that the higher transactivation potential of modified Gal4s show toxicity in mosquitoes when expressed from the oenocyte enhancer, presumably through a ‘squelching’ process (Gill and Ptashne 1988). The efficiency of precise cassette exchange with the A14 and A11 lines would appear to be at least 1 in 50 to 100 embryos injected, and thus similar to *piggyBac* transformation rates in this lab (Lynd and Lycett 2012; Lycett et al. 2012; Lombardo et al. 2009)

By assaying luciferase expression in crosses between a UAS responder line and alternative RCME lines that had the Gal4 cassette inserted in opposite orientations with respect to the *piggyBac* arms, we observed that relative activation potential was significantly higher when the Gal4 was inserted nearer to the right *piggyBac* arm (i.e. Orientations A > B Fig S3) in both larvae and adults. However, significant activation was detected in both orientations, as would be predicted from a local enhancer that acts upstream or downstream of a target gene (Khoury and Gruss 1983).

By fluorescently tagging both larval and adult oenocytes, we are able to visually follow the development of adult oenocytes during the 4^th^ instar to adult stage, with the parallel loss of larval oenocytes during late pupation. The timing of the mosquito oenocyte cell development pathways has been documented as early as 1960 (Christophers 1960), but generating a Gal4 driver for these cells has opened the possibility for a variety of cellular and functional analysis. For example, this tagging will enable isolation of pure cells for ‘omics’ analysis; however, initially we have used the oenocyte driver to examine the *in vivo* function of two CYP4G P450s annotated in *An. gambiae*.

Our previous *in vitro* analysis could only detect aldehyde decarbonylase activity for recombinant CYP4G16 when assayed with a C18 substrate, yet both genes are highly expressed in pupal and adult oenocytes, implicating both for roles in this cell type (Ingham et al. 2014; Balabanidou et al. 2016; Kefi et al. 2019). Furthermore, here extending immunolocalisation analysis of the abdomen integument to an earlier stage shows that both genes are also expressed in 4^th^ instar larval oenocytes, yet qPCR shows distinct temporal transcription patterns. *cyp4g16* is highly constitutively expressed throughout all life stages examined, and only increases 10-fold during 4^th^ instar and pupal stages compared to the 3^rd^ instar, whereas *cyp4g17* is barely detectable in 3^rd^ instar larvae, but is very highly induced in 4^th^ instar and late pupal stages (up to 10,000-fold). The temporal profiles would suggest that the two genes have distinct roles in *An. gambiae* related to life cycle stage, which correlates with the early and late onset of lethal pupal phenotypes observed in the RNAi knockdowns. It may also indicate that the expression of the two genes are regulated by different effectors. The regulation of cyp4G17 in 4^th^ instar larvae and pupae would suggest control linked to larval to adult transition, perhaps hormonal.

As female *An. gambiae* age there a relative decrease in abundance of low molecular weight (LMW) CHCs (up to heptacosane) and an increase in the relative abundance of higher MW CHCs including heptacosane, nonacosane, hentriacontane and methyl-hentriacontane (Caputo et al. 2005), which is recapitulated in the CHC analysis performed on pupal and adult stages here, and most clearly seen in the controls samples in Fig S10. When examining total CHC content in early pupae and surviving one day old adults it was clear that silencing either gene results in a similar 80% and 50% reduction respectively. Most strikingly, both gene knockouts have a significant reduction in abundance of nearly all CHCs at the pupal stage compared with the control, however no significant difference in abundance of individual CHCs between the two knockdowns were detected at this stage. Which would indicate both enzymes have major roles in CHC synthesis *in vivo* and have significant overlapping substrate specificity. The fact that both individual knockdowns produce 80% reductions in total pupal CHC also suggests a degree of synergism in CYP4G activity in 4^th^ instar larvae and pupae.

Significant differences in the two knockdowns were observed however when analysing relative abundance of each CHC compared with the total CHC content. The 16i knockdowns pupa had relatively less of the major HMW alkanes and methyl-alkanes compared with 17i knockdowns, whereas 17i had significantly lower ratios of the minor even-chain CHCs. The lower proportion of higher molecular weight CHCs would suggest that the earlier larval expression of *cyp4G16* is critical for generating the appropriate relative abundance of CHCs for early pupal survival, and the relative ratio of CHCs may be a contributing factor to the death observed in the early 16i pupae.

The CHC profiles from adults have to be addressed with the caveat that since the adults we analysed are outliers which have survived eclosion, and may well have less extreme changes than those which die and were excluded. Despite this, in the adult survivors, there is evidence of small differences in absolute CHC content between the knockdowns, in that *cyp4g16* knockdown adults show reduced levels of nonadecane compared with 17i individuals, whilst *cyp4g17* knockdowns show reduced levels of methyl-hentriacontane compared to *cyp4g16* knockdowns. The differences observed between 16i and 17i in relative proportion of individual CHC compared to total CHC were less pronounced in the adult than in the pupae, which may be a reflection of outlier analysis. In adults, nonadecane and nonacosane were reduced in 16i and, opposite to the pupal data, the HMW methyl-CHCs were relatively reduced in 17i compared to 16i. Indicative of an enhanced role for CYP4G17 in generating methyl-CHCs late in pupal development that perhaps facilitates the eclosion process.

The CHC profiling would thus suggest that the two enzymes have significant overlap of substrate specificity, and at the early pupal stage the enzymes work synergistically. At this stage, the CYP4G16 has been active since at least 3^rd^ instar and has slight dominance in the production of HMW CHCs. During later pupal to adult development, CYP4G17 appears dominate in the production of methyl branched chains. A propensity to generate methyl-CHCs was also observed when *cyp4g17* is ectopically expressed in cyp4G1 silenced in Drosophila (Kefi et al, 2019). It should be noted however that the temporal analysis is complicated by the unknown rate of deposition of internally stored CHCs to the cuticle. Furthermore, the effect of observed changes in sub-cellular localisation of the CYP4Gs at different developmental stages may play a role in enzyme activity. At the 4^th^ larval stage, we have previously shown both enzymes are localized in the periphery of the oenocyte cell membrane (Kefi et al. 2019), where synergistic function may be facilitated. In the developing adult oenocytes cyp4G16 maintains its peripheral cell localisation, whereas cyp4G17 was observed to localise more broadly throughout the endoplasmic reticulum (Kefi et al. 2019), which may alter which substrates the two enzymes encounter and their subsequent activities.

All insects possess one or more *cyp4g* P450s (phylogenetic tree in (Calla et al. 2018). In *D. melanogaster* their two annotated genes have distinct tissue specific expression patterns, although only *DmCyp4g1* is expressed in oenocytes throughout life and has a critical role in CHC production (Qui et al. 2012), whereas *cyp4g15* appears predominantly neuronal and of ill-defined function (Maibeche-Coisne et al. 2000). Oenocyte-specific *Dmcyp4g1* RNAi knockdown produces a temporally similar, partially incomplete, lethal pharate adult phenotype similar to that observed in *cyp4g17* knockdown mosquitoes. Although the mortality during eclosion of the vast majority of 17i mosquitoes precluded analysis of desiccation tolerance, the most parsimonious explanation for pharate adult death is severe desiccation intolerance. However, we cannot rule out other developmental irregularities, including a direct or indirect role for CHCs to facilitate escape from the pupal cuticle (Chiang et al. 2016). The more weakly penetrant lethal phenotype in *cyp4g16* knockdown pupae, allowed the demonstration of significant sensitivity to desiccation in surviving adults. This supports the hypothesis that a primary determinant of desiccation tolerance in *Anopheles* mosquitoes is total CHC content (Arcaz et al. 2016). This later study also demonstrated a positive correlation between octacosane and desiccation tolerance that was significantly depleted in the *cyp4g17* knockdown mosquitoes.

Transient knockdown of the single *cyp4gs* in holometabolous locusts (Yu et al. 2016) and pea aphids (Chen et al. 2016) also produce a desiccation sensitive phenotype. In the former case, the lethal phenotype could be rescued by maintenance at 90% humidity. However, attempts to rescue *cyp4g17* knockdown mosquitoes was unsuccessful when maintained at close to 100% during eclosion (not shown).

Although this is the first time that two *cyp4g*s have both been shown to be essential for CHC production in an insect, *in vitro* activity of recombinant proteins from the two annotated *cyp4g*s from the Mountain Pine beetle both convert long chain alcohols and aldehydes into HCs (MacLean et al. 2018). This led to speculation that one or both may be involved in CHC and pheromone production. However, their *in vivo* function and anatomical distribution is yet to be explored in this beetle. In this context, the finding that the single CYP4G from honey bees, that also decarbonylates aldehydes (Calla et al. 2018), is located in antennae and other chemosensory organs (Mao et al. 2015; Calla et al. 2018), as well as abdominal tergites associated with wax production, may suggest a second role in chemoreception for this P450 class. Further work will be needed in Anopheles mosquitoes to examine the functional role of *cyp4g* expression in sensory organs, since data from RNA-Seq (Pitts et al. 2011), indicate significant transcription of both *cyp4g16* and *cyp4g17* in *An. gambiae* antennae.

In summary, successful enhancer trapping of oenocyte regulatory regions by RMCE, and the subsequent use of stable RNAi have enabled us to examine the role of two highly expressed CYPs in CHC production and paved the way for in depth analysis of the metabolic roles of oenocytes in mosquitoes. For example, the driver line will be critical for oenocyte-specific gene knockdown and ectopic over-expression to examine the complex metabolic functions of these cells in the malaria mosquito, and to explore their potential role in pheromone production, mating behaviour and longevity (Joseph et al. 2018; Combs et al. 2018).

## Supporting information

primer sets

Summary of RCME experiments performed with different Gal4 constructs in A14 and A11 docking lines.

pupal lethality

GCMS annotation

Statistical analysis of CHC normalized to Internal 363 standard

Statistical analysis of CHC normalized to Total CHC content

Schematic of the constructs used for transformation

Summary of generation of docking lines

Phenotypic characterisation of RCME lines

Dissection of Female Adult and Pupa derived from A14Gal4 x UAS-mcherry cross

Images of A11Gal4 X UAS-nlsYFP progeny

CYPG16 and CYP4G17 larval expression

Schematic of the constructs used for gene knockout

Western analysis of 16i,17i and A14A1 early pupae

Plots of relative abundance of CHCs with respect to internal standard

Plots of relative abundance of CHCs with respect to total CHC

## Acknowledgements

We’d like to thank Amalia Anthousi for great technical assistance in mosquito husbandry and Dolphine Amenya for her major role in the initial generation and characterisation of the pBac-CFP lines. We also thanks Jacob Chapman and Judit Bagi for assistance with qRT-PCR analysis. We also are indebted to Mark Prescott and Rob Beynon who performed the CHC analysis at the Centre for Proteome Research, University of Liverpool, Mike Povelones for providing LRIM1 clones and antibodies, Leonie Ringrose for the PKC40 plasmid, and to Hilary Ranson for thoughtful comments on the manuscript. We are also extremely grateful for two anonymous reviewers that greatly improved the manuscript. The work was partly funded by the EU FP7 Avecnet Grant ID: 265660.

## Notes

#### Summary of Updates

Figures and legends have been improved.

