## Supplementary material for "Development of a functional genetic tool for Anopheles gambiae oenocyte characterisation: application to cuticular hydrocarbon synthesis": primer sets

Table S1

| | primer name | sequence 5'-3' |
|---|---|---|
| **general cloning primers** | | |
| | BGH-F | ggatccggcccgtttaaacccgctg |
| | BGH-R | gcggccgcaagccatagagcccaccgc |
| | attP-F | ctcgagcgaagccccggcggcaac |
| | attP-R | gcggccgccgcgctcgcgcgactgac |
| | LRIM1-F | ggatccatgacgacgacaagatggc |
| | LRIM1-R | ggatcctctagattggttactgggcg |
| | attPPSTI-F | ctgcagcgaagccccggcggcaac |
| | attPPSTI-R | ctgcagcgcgctcgcgcgactgac |
| | HSP-F | ttgtgcggccgccgagcgccggagtataaat |
| | HSP-R | ttgtgaattctccctattcagagttctcttcttg |
| | attB5-F | ttctcttaaggtcgacgatgtaggtcacg |
| | attB5-R | ttctactagtcactatagggcgaattgggta |
| | attB3-F | ttctagatctgtcgacgatgtaggtcacg |
| | attB3-R | ttctatgcatcactatagggcgaattgggta |
| | Red-F | tcttgctagccgcacggttcccacaat |
| | Red-R | ttctatgcatttaagatacattgatgagtttggac |
| | gyp1F | tgttaagcttatgcatcaatgtatcttaactactcacgt |
| | gyp1R | tgttgcatgcactagaattgatcggctaaat |
| | gyp2F | tgttggatcccaatgtatcttaactactcacgt |
| | gyp2R | tgttggatccgcggccgcactagaattgatcggctaaat |
| | RINcoF | ttctgaattcttctcagatctttctcgatatcttctcgctagcttctcctcgagttctcccatggttct |
| | RINcoR | agaaccatgggagaactcgaggagaagctagcgagaagatatcgagaaagatctgagaagaattcagaa |
| | Link2F | ttctgcggccgctcttcctatgcattctt |
| | Link2R | aagaatgcataggaagagcggccgcagaa |
| **hairpin construction** | | |
| **Cyp16RNAi-gDNA** | Cyp16RNAi-F | aacagctagcccgtgttccgcacaatta |
| | Cyp16 bridge | ggactacttataagcaccgttgacgccatttgtgtc |
| **Cyp17RNAi-gDNA** | Cyp17RNAi-1 | aacagctagctaccgggcccggtcgcct |
| | Cyp17RNAi-2 | aatgtgtgacatcctgcactgcaagaaagtttaaatttg |
| **Cyp17RNAi-cDNA** | Cyp17RNAi-3 | caaatttaaactttcttgcagtgcaggatgtcacacatt |
| | Cyp17RNAi-4 | aacaccatggaacacctacgtacccactgatcgggaac |
| **Cyp17RNAi-fusion** | Cyp17RNAi-1 | aacagctagctaccgggcccggtcgcct |
| | Cyp17RNAi-5 | aacaccatggaacacctacg |
| **Orientation of RMCE** | | |
| | pBAC right | tttgcctttcgccttattttaga |
| | pBAC left | tgacgagcttgttggtgaggattct |
| | Internal -1 | cgagggttcgaaatcgataa |
| | Internal -2 | tcggtttttctttggagcac |
| | Internal -3 | ccccgtaatgcagaagaaga |
| | Internal -4 | aaggaaaaagctgcactgct |
| **Southern Probes** | | |
| **pBac-Left** | PBLFor | ggtcatctgtggacatgtgg |
| | PBLRev | gatgacgagcttgttggtga |
| **pBac-Right** | PBRFor | cgcatgtgttttatcggtct |
| | PBRev | aaaaggtccaaagtcgcaaa |
| **qRT-PCR Primers** | | |
| **Amplicon/efficiency** | | |
| **Cyp16-1b/99.7** | Cyp16-q1bF | tgcagaacgagaatgggaaagtc |
| | Cyp16-q1bR | actcccattgcagtttctagaag |
| **Cyp16-2a/99.6** | Cyp16-q2aF | gacaaatcccctgaataccgc |
| | Cyp16-q2aR | ccatttgtgtccggtgctta |
| **Cyp17-2a/97.6** | Cyp17-q2aF | gaaacgttcaacccggacaa |
| | Cyp17-q2aR | tactttcgacccacacagct |
| **Cyp17-2b/99.3** | Cyp17-q2bF | agctgtgtgggtcgaaagta |
| | Cyp17-q2bR | agtccttctcggtcaggttg |
| **RT-PCR Primers** | | |
| **AG003206RA** | 3206AF | gaggatgcggaacagggt |
| | 3206AR | caatttcactgtccgtcacca |
| **AG003206RB** | 3206BF | tgtgtgaggttcaataaggtgtg |
| | 3206BR | gcaatttcactgtccgtcgt |
| **AG003206RC** | 3206CF | tcactcttcacctgccgtag |
| | 3206CR | caatttcactgtccgtccgg |
| **S7** | S7FOR | gtagctgctgcaaacttcgg |
| | S7REV | ggcgatcatcatctacgt |
