## Supplementary material for "Development of a functional genetic tool for Anopheles gambiae oenocyte characterisation: application to cuticular hydrocarbon synthesis": Summary of RCME experiments performed with different Gal4 constructs in A14 and A11 docking lines.

Table S2 Summary of RCME experiments performed with different Gal4 constructs in A14 and A11 docking lines.

| A14 | Gal4 form | Eggs injected/larvae hatched | No of Adult Pools giving dsred positive larvae/total pools | Dsred only positive/ Dsred and CFP F1 larvae | Isofemale lines produced | Insert Orientation |
|---|---|---|---|---|---|---|
| | | | | | Gth | A |
| | | | | | Ell | A |
| | | | | | Esm | C |
| | Gal4 | 406/133 | 4/8 | 105/50 | Erm | B |
| | | | | | Alx | F |
| | | | | | Ezb | D |
| | | | | | Erc | D + |
| | Gal4Δ | 472/214 | 7/9 | 26/18 | All died | |
| | GFY | 578/187 | 6/12 | 44/39 | All died | |
| | FF | 389/91 | 3/4 | 3/3 | 1 dsred/cfp survivor | E |
| A11 | Gal4 | 638/292 | 9/14 | 164/20 | Kti | A |
| | | | | | Dph | A |
| | | | | | Nbl | B |
| | | | | | Krk | B |
| | | | | | Ics | B + |
| | | | | | Npt | C |

Table indicates the four different versions of Gal4 transactivator used for RCME in the A14 docking line, and the use of native Gal4 for RCME in A11.
Also indicated are the number of eggs injected and larval hatch numbers for each construct. Total number of adult G0 pools set up for each construct and number of pools giving rise to transgenic progeny.
Isofemale lines produced refers to lines analysed for insert orientation, letters A to F refer to the orientation of the attB cassette relative to piggyBac arms (see Figure S3)
