## Supplementary material for "Development of a functional genetic tool for Anopheles gambiae oenocyte characterisation: application to cuticular hydrocarbon synthesis": pupal lethality

| | Surviving adults | Pupae early death/drowned | Late pupae/partially eclosed adults |
|---|---|---|---|
| A11 homo | 93 | 4 | 4 |
| A14 controls | 122 | 2 | 1 |
| 16i | 39 | 134 | 39 |
| 17i | 8 | 0 | 81 |
| | proportion of surviving adults | proportion of Pupae early death/drowned | proportion of Late pupae/partially eclosed adults |
| A11 homo | 0.92 | 0.04 | 0.04 |
| A14 controls | 0.98 | 0.02 | 0.01 |
| 16i | 0.18 | 0.63 | 0.18 |
| 17i | 0.09 | 0.00 | 0.91 |

**Table S3** Typical scoring of parental or cross larval progeny for stage of death when grown at low density in 30x30x5cm tray.

A11 homo are homozygous parental docking strain generated through crossing of A11 docking strain (cfp) with another A11 recipient strain (UAS:P3) (yfp) thus having different markers at same locus, crossing those transheterozygous progeny having both markers and rearing the only cfp marked progeny which are homozygous at that locus.

A14 controls are siblings from A14Gal x 16i crosses that are heterozygous for the A14Gal4 allele (cfp) and lack the UAS hairpin allele (yfp)

16i and 17i are progeny from respective A14Gal crosses with UA S-16i or 17i and carry both selectable markers

Surviving adults are those which eclose and are free flying,

pupae early death/drowned are those pupae that sink to the bottom of the water and die,

late pupae are those that are pharate adults that remain floating or are partially eclosed adults
