## Supplementary material for "Development of a functional genetic tool for Anopheles gambiae oenocyte characterisation: application to cuticular hydrocarbon synthesis": GCMS annotation

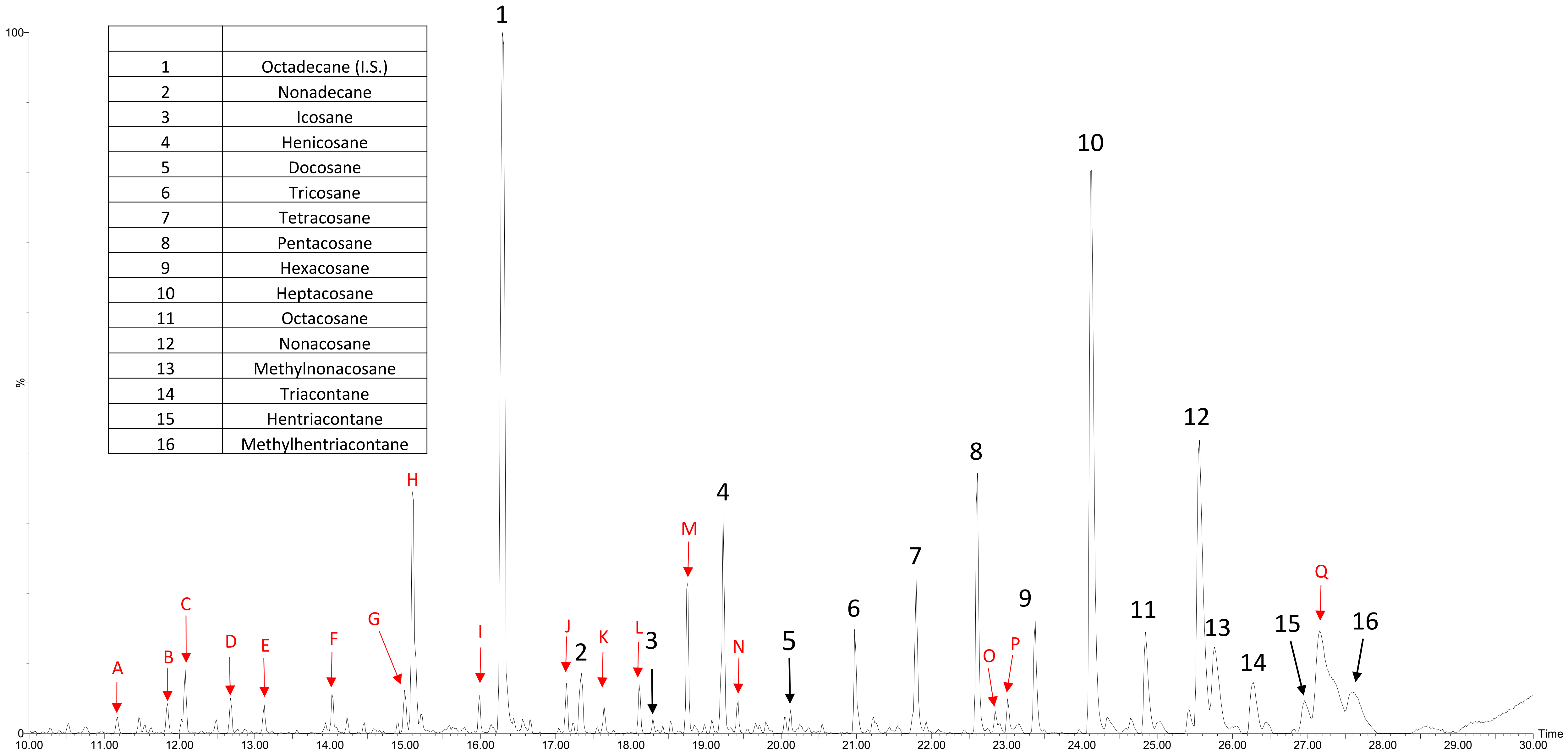


| | |
|---|---|
| 1 | Octadecane (I.S.) |
| 2 | Nonadecane |
| 3 | Icosane |
| 4 | Henicosane |
| 5 | Docosane |
| 6 | Tricosane |
| 7 | Tetracosane |
| 8 | Pentacosane |
| 9 | Hexacosane |
| 10 | Heptacosane |
| 11 | Octacosane |
| 12 | Nonacosane |
| 13 | Methylnonacosane |
| 14 | Triacontane |
| 15 | Hentriacontane |
| 16 | Methylhentriacontane |

| Peak | Potential identities from NIST Library search |
|---|---|
| A | No match |
| B | No match |
| C | Siloxane |
| D | Ethopropazine |
| E | No match |
| F | No match |
| G | ester? |
| H | 5-methyl-2-N,N-dimethyl-aminobenzophenone hydrazone |
| I | 3-methyl-heptadecane |
| J | 1,2-benzenedicarboxylic acid,butyl-2-methyl-propyl ester |
| K | No match |
| L | Bis(2-methoxyethyl)pthalate |
| M | 1-cyclohexene-1-carboxylic acid,4-(1,5-dimethyl-3-oxohexyl)methylester |
| N | methyl ester? |
| O | 2(1H)-isoquinoline |
| P | Carboxamide, 1-cyano-N,N-diphenyl |
| Q | Cholesterol |
