## Supplementary material for "Development of a functional genetic tool for Anopheles gambiae oenocyte characterisation: application to cuticular hydrocarbon synthesis": Statistical analysis of CHC normalized to Internal 363 standard

File S2

| Pupae | | | | | | | | | |
|---|---|---|---|---|---|---|---|---|---|
| | Df | Sum Sq | Mean Sq | F value | Pr(>F) | adjusted | 16i-A14Gal4 | 17i-A14Gal4 | 17i-16i |
| **Nonadecane** | 2 | 5.74E-06 | 2.87E-06 | 0.628130557 | 0.541205786 | 0.541205786 | 0.546886958 | 0.975424444 | 0.677705552 |
| **Icosane** | 2 | 4.39E-06 | 2.19E-06 | 4.186287441 | 0.026086193 | 0.034781591 | 0.075038815 | 0.032223794 | 0.918578949 |
| **Henicosane** | 2 | 1.27361778 | 0.63680889 | 295.5496359 | 4.41E-19 | 1.06E-17 | 4.22E-15 | 4.22E-15 | 0.988773226 |
| **Docosane** | 2 | 0.001346614 | 0.000673307 | 30.63524517 | 1.13E-07 | 3.03E-07 | 3.78E-07 | 2.05E-06 | 0.787350475 |
| **Tricosane** | 2 | 0.527628274 | 0.263814137 | 129.229084 | 1.49E-14 | 8.95E-14 | 2.82E-13 | 1.97E-13 | 0.975534411 |
| **Tetracosane** | 2 | 0.002507797 | 0.001253898 | 25.533573 | 5.96E-07 | 1.19E-06 | 6.95E-06 | 2.19E-06 | 0.898232584 |
| **Pentacosane** | 2 | 0.711341717 | 0.355670858 | 169.1877755 | 5.33E-16 | 6.39E-15 | 1.68E-14 | 1.09E-14 | 0.898929975 |
| **Hexacosane** | 2 | 0.007604956 | 0.003802478 | 22.08164778 | 2.08E-06 | 3.84E-06 | 2.51E-05 | 6.15E-06 | 0.856683735 |
| **Heptacosane** | 2 | 0.837688895 | 0.418844447 | 141.0587751 | 5.09E-15 | 4.07E-14 | 3.26E-14 | 2.84E-13 | 0.36876591 |
| **Octacosane** | 2 | 0.002522945 | 0.001261473 | 8.391911006 | 0.001464456 | 0.002343129 | 0.015971024 | 0.001523349 | 0.62024471 |
| **Nonacosane** | 2 | 0.05378547 | 0.026892735 | 42.3704322 | 4.7E-09 | 1.61E-08 | 1.69E-08 | 1.29E-07 | 0.681026615 |
| **Methylnonacosane** | 2 | 0.012134312 | 0.006067156 | 16.97945329 | 1.68E-05 | 2.88E-05 | 3.97E-05 | 0.000171282 | 0.848430909 |
| **Triacontane** | 2 | 0.000449098 | 0.000224549 | 4.010876394 | 0.02984348 | 0.037697027 | 0.56005366 | 0.024430392 | 0.200935574 |
| **Hentriacontane** | 2 | 0.000548886 | 0.000274443 | 5.386937121 | 0.010747938 | 0.016121907 | 0.076274138 | 0.00982603 | 0.640328955 |
| **Methylhentriacontane** | 2 | 0.023423543 | 0.011711771 | 38.6838602 | 1.18E-08 | 3.55E-08 | 5.95E-08 | 1.8E-07 | 0.893995197 |

| Adult | | | | | | | | | |
|---|---|---|---|---|---|---|---|---|---|
| | Df | Sum Sq | Mean Sq | F value | Pr(>F) | adjusted | 16i-A14Gal4 | 17i-A14Gal4 | 17i-16i |
| **Nonadecane** | 2 | 0.002196777 | 0.001098389 | 3.906145801 | 0.032360897 | 0.051777434 | 0.114296835 | 0.82890949 | 0.033666585 |
| **Icosane** | 2 | 7.71E-05 | 3.86E-05 | 2.202690131 | 0.129973872 | 0.189052904 | 0.210735872 | 0.160888643 | 0.986872327 |
| **Henicosane** | 2 | 0.011487872 | 0.005743936 | 4.87295357 | 0.015598746 | 0.031197491 | 0.214317018 | 0.011634114 | 0.359819541 |
| **Docosane** | 2 | 7.17E-06 | 3.59E-06 | 0.408293152 | 0.66881975 | 0.764365429 | 0.999992932 | 0.717808457 | 0.715653739 |
| **Tricosane** | 2 | 0.002573282 | 0.001286641 | 1.824307188 | 0.180661 | 0.240881334 | 0.948013987 | 0.317824607 | 0.192676047 |
| **Tetracosane** | 2 | 0.000159392 | 7.97E-05 | 0.110431421 | 0.895850082 | 0.930281392 | 0.967818776 | 0.972238582 | 0.885940194 |
| **Pentacosane** | 2 | 0.046707124 | 0.023353562 | 6.073374216 | 0.00663772 | 0.017700587 | 0.025568334 | 0.009267185 | 0.904187363 |
| **Hexacosane** | 2 | 0.007200956 | 0.003600478 | 10.4928153 | 0.00042502 | 0.001700081 | 0.013614282 | 0.000347248 | 0.337861749 |
| **Heptacosane** | 2 | 1.744509209 | 0.872254605 | 59.74467245 | 1.22E-10 | 1.95E-09 | 1.37E-09 | 1.32E-09 | 0.999836657 |
| **Octacosane** | 2 | 0.005609191 | 0.002804596 | 4.357821633 | 0.022899257 | 0.040709789 | 0.234241218 | 0.017557751 | 0.423046143 |
| **Nonacosane** | 2 | 0.506855616 | 0.253427808 | 44.97774232 | 2.54E-09 | 2.03E-08 | 1.04E-08 | 6.36E-08 | 0.728601001 |
| **Methylnonacosane** | 2 | 0.01987656 | 0.00993828 | 8.239507576 | 0.001609292 | 0.005149735 | 0.117390607 | 0.001068285 | 0.131715992 |
| **Triacontane** | 2 | 0.000536265 | 0.000268133 | 0.82848987 | 0.447507369 | 0.5507783 | 0.932448946 | 0.435283452 | 0.649399241 |
| **Hentriacontane** | 2 | 0.001541524 | 0.000770762 | 5.231021732 | 0.012020598 | 0.027475652 | 0.031093547 | 0.019593004 | 0.977932676 |
| **Methylhentriacontane** | 2 | 0.05175455 | 0.025877275 | 14.21048634 | 6.08E-05 | 0.000324197 | 0.069453603 | 3.77E-05 | 0.015592099 |

**Statistical analysis performed by ANOVA with Benjamini-Hochber p-value adjustment followed by Tukey's HSD (Honestly Significant Difference) for pairwise comparisons.**
