## Supplementary material for "Development of a functional genetic tool for Anopheles gambiae oenocyte characterisation: application to cuticular hydrocarbon synthesis": Statistical analysis of CHC normalized to Total CHC content

File S3

| Pupae | | | | | | | | | |
|---|---|---|---|---|---|---|---|---|---|
| | Df | Sum Sq | Mean Sq | F value | Pr(>F) | adjusted | 16i-A14Gal4 | 17i-A14Gal4 | 17i-16i |
| Nonadecane | 2 | 0.002964404 | 0.001482202 | 19.52189658 | 5.7E-06 | 1.37E-05 | 6.04E-06 | 0.000345131 | 0.296713262 |
| Icosane | 2 | 4.61E-05 | 2.31E-05 | 8.688041127 | 0.001221518 | 0.001724496 | 0.000997878 | 0.026833768 | 0.393850303 |
| Henicosane | 2 | 0.10662037 | 0.053310185 | 21.26583763 | 2.84E-06 | 8.53E-06 | 1.49E-05 | 1.65E-05 | 0.999090461 |
| Docosane | 2 | 0.000848895 | 0.000424447 | 6.140928965 | 0.006336049 | 0.007241199 | 0.01094457 | 0.018695332 | 0.972575685 |
| Tricosane | 2 | 0.023592006 | 0.011796003 | 10.68330849 | 0.000381983 | 0.000572974 | 0.00150066 | 0.0010298 | 0.988599384 |
| Tetracosane | 2 | 0.002306253 | 0.001153126 | 19.65944833 | 5.39E-06 | 1.37E-05 | 4.51E-06 | 0.00066411 | 0.163687608 |
| Pentacosane | 2 | 0.019462146 | 0.009731073 | 12.85712819 | 0.000119504 | 0.000204864 | 0.002896797 | 0.000126546 | 0.469307648 |
| Hexacosane | 2 | 0.006844289 | 0.003422145 | 23.07255045 | 1.44E-06 | 5.74E-06 | 1.09E-06 | 0.000374153 | 0.08731959 |
| Heptacosane | 2 | 0.049217505 | 0.024608752 | 91.68379307 | 9.19E-13 | 2.2E-11 | 1.26E-12 | 0.012951595 | 5.39E-10 |
| Octacosane | 2 | 0.014456187 | 0.007228094 | 29.43623919 | 1.65E-07 | 7.9E-07 | 1.19E-07 | 9.86E-05 | 0.039025999 |
| Nonacosane | 2 | 0.004703444 | 0.002351722 | 11.99649614 | 0.000187076 | 0.000299322 | 0.105845244 | 0.000120044 | 0.026246013 |
| Methylnonacosane | 2 | 0.001028599 | 0.000514299 | 8.57752533 | 0.001306699 | 0.001742265 | 0.001122434 | 0.445655561 | 0.023973277 |
| Triacontane | 2 | 0.008399461 | 0.004199731 | 37.12538979 | 1.78E-08 | 2.14E-07 | 1.02E-08 | 7.91E-05 | 0.004147034 |
| Hentriacontane | 2 | 0.005140714 | 0.002570357 | 31.59135943 | 8.5E-08 | 5.1E-07 | 6.86E-08 | 3.74E-05 | 0.052849564 |
| Methylhentriacontane | 2 | 0.001462575 | 0.000731288 | 18.34283159 | 9.31E-06 | 2.01E-05 | 5.62E-06 | 0.00515732 | 0.039269438 |
| Adult | | | | | | | | | |
| | Df | Sum Sq | Mean Sq | F value | Pr(>F) | adjusted | 16i-A14Gal4 | 17i-A14Gal4 | 17i-16i |
| Nonadecane | 2 | 0.003924856 | 0.001962428 | 13.74466154 | 7.64E-05 | 0.000203787 | 0.985187966 | 0.000242075 | 0.000374991 |
| Icosane | 2 | 4.72E-05 | 2.36E-05 | 2.086551188 | 0.143674756 | 0.164199721 | 0.356056278 | 0.134257156 | 0.826090481 |
| Henicosane | 2 | 0.008623884 | 0.004311942 | 11.91868394 | 0.000194957 | 0.000389915 | 0.000306687 | 0.001894252 | 0.76984317 |
| Docosane | 2 | 0.000173295 | 8.66E-05 | 33.00163898 | 5.61E-08 | 2.99E-07 | 2.95E-07 | 6.4E-07 | 0.949347834 |
| Tricosane | 2 | 0.009651175 | 0.004825587 | 18.8525287 | 7.51E-06 | 3E-05 | 5.46E-06 | 0.001443165 | 0.107924731 |
| Tetracosane | 2 | 0.02668941 | 0.013344705 | 34.00948679 | 4.2E-08 | 2.99E-07 | 8.42E-06 | 4.52E-08 | 0.116408304 |
| Pentacosane | 2 | 0.011129425 | 0.005564713 | 5.620536009 | 0.009104478 | 0.014567164 | 0.015336847 | 0.024720837 | 0.977243079 |
| Hexacosane | 2 | 0.000533876 | 0.000266938 | 3.872034387 | 0.033229338 | 0.044305784 | 0.054971825 | 0.998565413 | 0.061234656 |
| Heptacosane | 2 | 0.15794855 | 0.078974275 | 59.89097595 | 1.18E-10 | 1.89E-09 | 3.34E-10 | 7.85E-09 | 0.344865467 |
| Octacosane | 2 | 0.000581486 | 0.000290743 | 1.764855566 | 0.190394515 | 0.203087482 | 0.611873146 | 0.629395722 | 0.164240672 |
| Nonacosane | 2 | 0.051011106 | 0.025505553 | 12.82829714 | 0.000121283 | 0.000277217 | 8.17E-05 | 0.136714531 | 0.013684542 |
| Methylnonacosane | 2 | 0.002097027 | 0.001048513 | 2.890518965 | 0.072862549 | 0.089676984 | 0.65123689 | 0.311644762 | 0.061848537 |
| Triacontane | 2 | 0.00187331 | 0.000936655 | 5.922326646 | 0.007369516 | 0.013101361 | 0.005218792 | 0.213313373 | 0.218595103 |
| Hentriacontane | 2 | 9.12E-05 | 4.56E-05 | 0.316281037 | 0.731518364 | 0.731518364 | 0.721783792 | 0.849907969 | 0.971477337 |
| Methylhentriacontane | 2 | 0.015039133 | 0.007519566 | 14.4858997 | 5.32E-05 | 0.000170206 | 0.026434722 | 0.037174544 | 3.18E-05 |
| Statistical analysis performed by ANOVA with Benjamini-Hochber p-value adjustment followed by Tukey's HSD (Honestly Significant Difference) for pairwise comparisons. | | | | | | | | | |
