## Supplementary material for "Development of a functional genetic tool for Anopheles gambiae oenocyte characterisation: application to cuticular hydrocarbon synthesis": Schematic of the constructs used for transformation

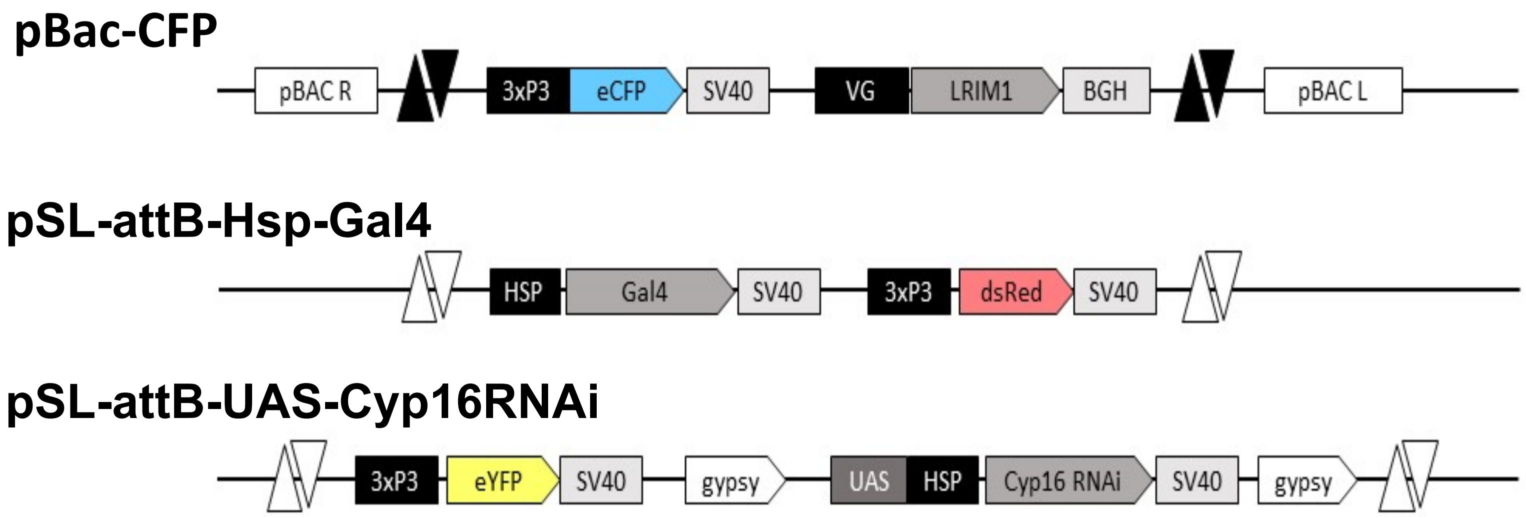


**Figure S1 Schematic of the constructs used for transformation**

pBac-CFP [attP-LRIM1], pSL-attB-Hsp-Gal4[3xP3-dsRed] and pSL-attB-UAS-Cyp16RNAi cassettes. The pSL-attB-UAS-Cyp17RNAi cassette (not shown) is identical to pSL-attB-UAS-Cyp16RNAi except for the Cyp4g16RNAi cassette is replaced with the Cyp4g17RNAi cassette. Vg- An. gambiae Vitellogenin2 promoter, LRIM1- Leucine Repeat Immune Molecule 1 cDNA carrying a MYC tags, BGH-Bovine growth hormone terminator, SV40-SV40 termination sequences, gypsy- Drosophila Gypsy insulator element, Paired black triangles- attP sites, Paired white triangles, attB sites.
