## Supplementary material for "Development of a functional genetic tool for Anopheles gambiae oenocyte characterisation: application to cuticular hydrocarbon synthesis": Summary of generation of docking lines

A

| PBAC-CFP injections | Embryos Injected | $G_o$ Larvae Hatched | $G_o$Pupa survived | No of fluorescent $G_1$ larvae | Independent single copy lines from inverse PCR analysis/total isofemale lines generated |
|---|---|---|---|---|---|
| | **704** | **112** | **77** | **55** | **4/10** |

B

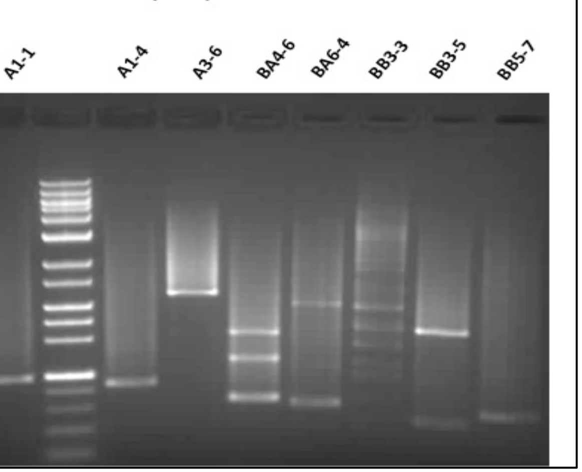


C

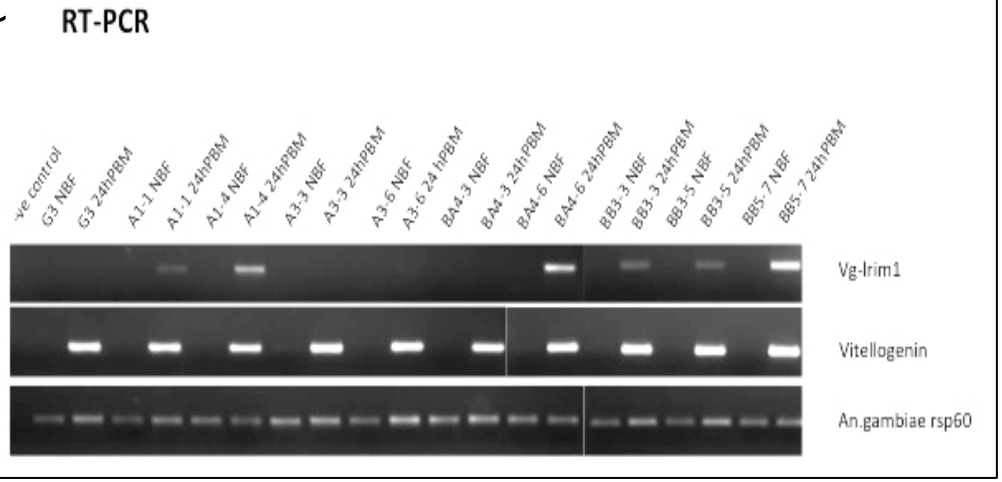


D

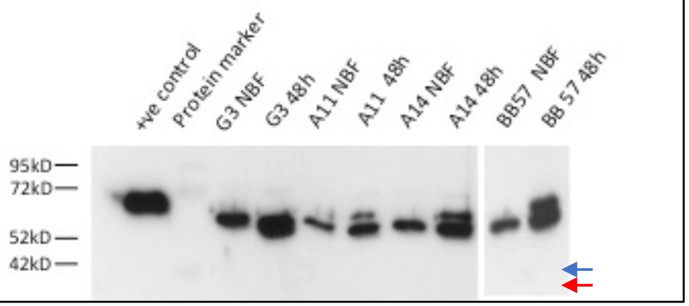


E

| Transgenic line | Insertion site | IPCR sequence | | Nearest annotated gene |
|---|---|---|---|---|
| A11 | 2R 33,858,877-80 | Left **TTAA**CATCCTCTTACTTCTAAAGTAAGACAAACACAT | Right **TTAA**ACAATACCCCAATTGGGTAGATTCAGTCTA | Within intron of AG003206 orthologue of CG13907 |
| A14 | 2R 48,392077-80 | Left: **TTAA**TTTAAATTTGAATTGAAATTCGTTCGGACGA | Right **TTAA**AAAGGAGCCGTTCGCTTTTGACGAGCAGACTATATTGTTCAG | 30 KB TO AGAP013096 CORTO ORTHOLOGUE 50KB TO CANOE ORTHOLOGUE NIETHER oenocyte restricted in drosophila |
| BB57 | 3L 38,793,391-94 | L **TTAA**GACCTCGATGAATTGAAGCTCATAAAATTGTT | Right **TTAA**AGGTCTCTTCAATTTACGTTAAACACTCCAAAATTC | 705 bp 5' to AGAP012209-RA No 1:1 drosophila orthologue |

F

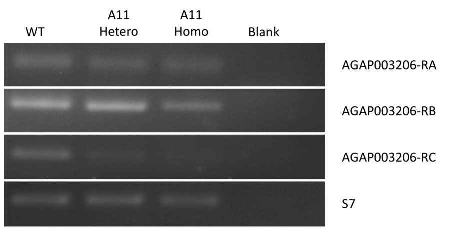


**Figure S2 Summary of generation of docking lines** A indicates summary data for the injection of embryos, screening of progeny and isofemale line production. B Inverse PCR to identify single insertions of docking construct from isofemale lines. C RT-PCR analysis to define expected inducible transcription of blood meal inducible LRIM1, controlled with Vg and ribosomal p60 specific primers, D Western analysis of LRIM1 induced expression in indicated single insertion isofemale lines (blue arrow – tagged LRIM, red arrow native LRIM1) using LRIM1 specific antibody. E Sequencing and identification of piggyBAC insertion sites in selected lines and indication of nearest PEST annotated genes to insertion site. F Semi quantitative RT-PCR of the three indicated, annotated isoforms of AG003206 (vectorbase) using exon-spanning primers (Sup Table3) under non saturating conditions (28 cycles) compared with housekeeping gene ribosomal protein S7 (22 cycles) for RNA isolated from heterozygous A11 (A11 Hetero), homozygous A11 (A11 homo), non transgenic siblings (WT) and water (blank) control).
