## Supplementary material for "Development of a functional genetic tool for Anopheles gambiae oenocyte characterisation: application to cuticular hydrocarbon synthesis": Phenotypic characterisation of RCME lines

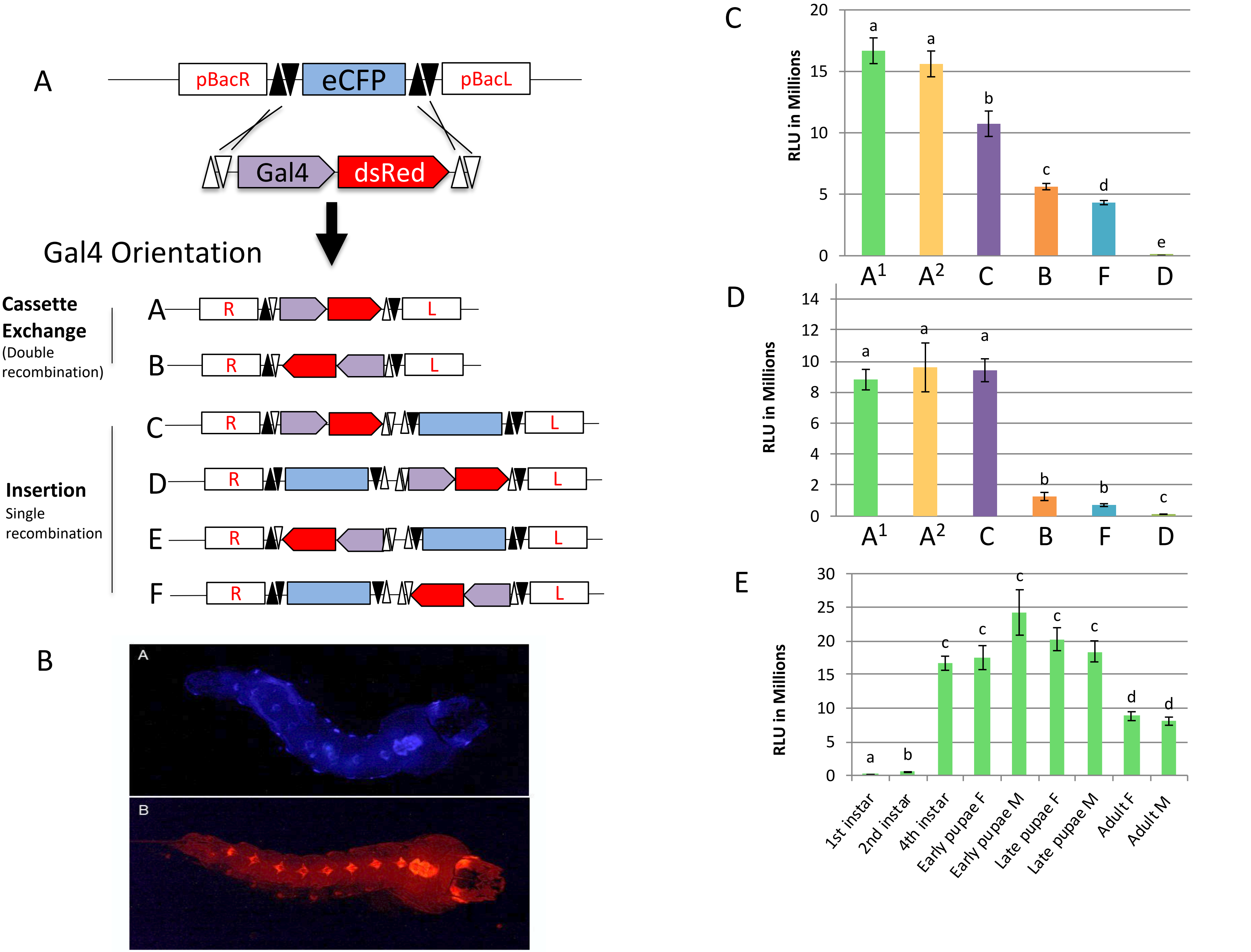
A
pBacR
eCFP
pBacL
Gal4
dsRed
Gal4 Orientation
Cassette Exchange
(Double recombination)
Insertion
Single recombination
A
B
C
D
E
F
R
L
B
A
B
C
RLU in Millions
20
15
10
5
0
a
a
b
c
d
e
A1
A2
C
B
F
D
D
RLU in Millions
14
12
10
8
6
4
2
0
a
a
a
b
b
c
A1
A2
C
B
F
D
E
RLU in Millions
30
25
20
15
10
5
0
a
b
c
c
c
c
c
d
d
1st instar
2nd instar
4th instar
Early pupae F
Early pupae M
Late pupae F
Late pupae M
Adult F
Adult M

**Figure S3 Phenotypic characterisation of RCME lines**

A Cartoon illustration of RCME process giving rise to six possible outcomes of transgene insertion, cassette exchange (in orientations A and B), or cassette integration (in orientations C-F). B: Images of 3$^{rd}$ instar larvae. top docking line A14 with pBac-CFP [attP-LRIM1] cassette showing eCFP expression in the neuronal tissue as directed by the 3Px3 promoter and in the oenocytes. bottom Line A14 after RMCE with the enhancer trapping cassette pSL-attB-Hsp-Gal4[3xP3-dsRed] showing dsRed expression in the same pattern as the A14 docking line. C and D: Mean luciferase activity in C: whole 4$^{th}$ instar larvae and D whole adult female for crosses between different orientation A14 Hsp-Gal4 driver lines (indicated along the X axis, see Table S1) and the UAS responder line, "Wnd". All mosquitoes were heterozygous for Gal4 and UAS cassettes (N = 14 minimum). RLU = Relative Light Units. Error bars show standard errors. Bars not sharing a letter differ significantly (Mann Whitney, $p<0.01$). E: Mean luciferase activity in various life stages of progeny from crosses between A14Gal4 (A1) driver line and the UAS responder line, Wnd. Life stage and sex are indicated along the X axis. All mosquitoes were heterozygous for Gal4 and UAS cassettes (N = 6). RLU = Relative Light Units. Error bars show standard errors. Significant differences are indicated by differing letters (Mann Whitney, $p<0.01$).

During the screening of F1 progeny from A14 RCME expts we observed a few larvae having noticeably greater dsRed expression than the majority, which was likely due to multiple copies of dsRed integrating into the genome. Although we localised one insertion to the D orientation, we could not verify the presence or define locations of other insertion/s, in this line, however,luciferase assays indicated higher levels in adult females in this line compared to all the other canonical transgenic lines obtained. A similar aberrant line displaying atypical dsRed fluorescence was also obtained during RCME of the neutral A11 line, suggesting that non-canonical transformation events may be a common occurrence during RCME.
