## Supplementary material for "Development of a functional genetic tool for Anopheles gambiae oenocyte characterisation: application to cuticular hydrocarbon synthesis": Dissection of Female Adult and Pupa derived from A14Gal4 x UAS-mcherry cross

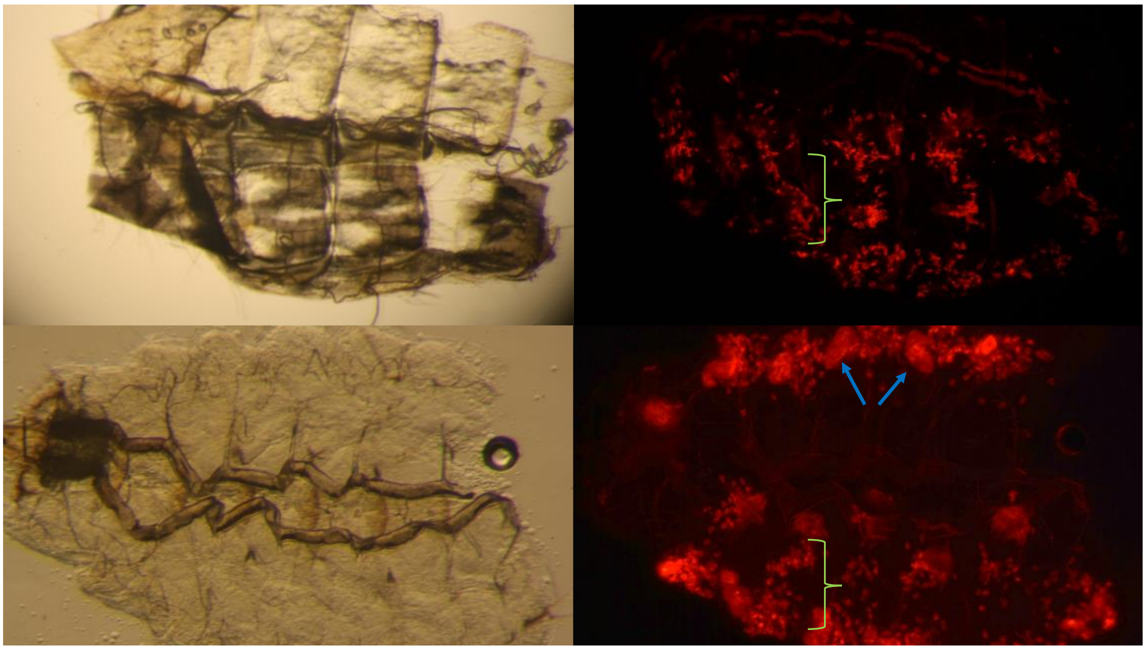

Figure S4: Dissection of Female Adult (Upper) and Pupa (Lower) derived from A14Gal4 x UAS-mcherry cross to flatten abdomen and expose dorsal and ventral surfaces without midgut. Left, brightfield images. Right, corresponding images taken through RFP filter. Brackets indicate adult type oenocyte cluster that develop in 4$^{th}$ instar larvae on ventral abdominal integument and are maintained throughout pupae and adult. Blue arrows indicate larval oenocytes remaining within the pupae.
