## Supplementary material for "Development of a functional genetic tool for Anopheles gambiae oenocyte characterisation: application to cuticular hydrocarbon synthesis": Images of A11Gal4 X UAS-nlsYFP progeny

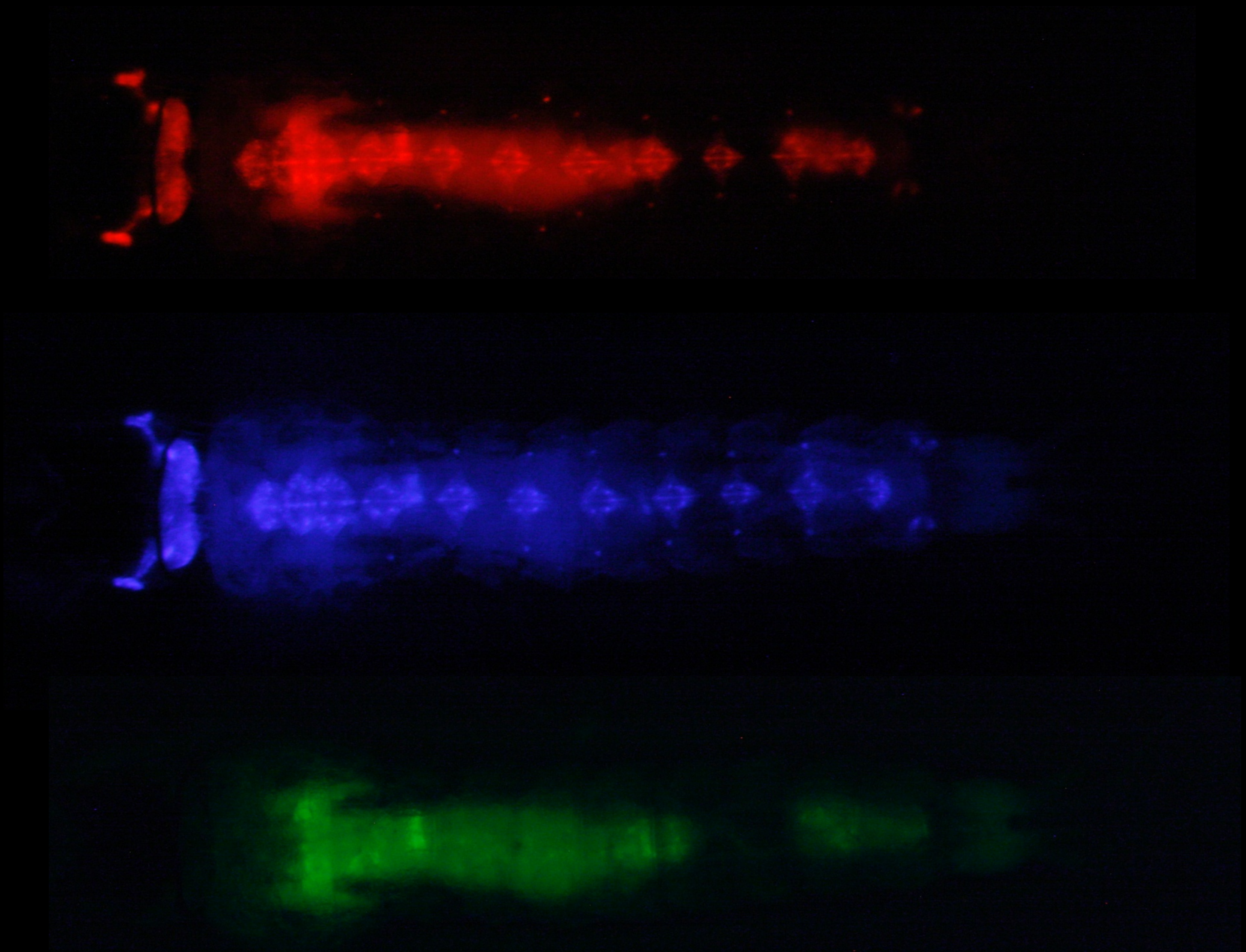

Figure S5, A11Gal4 X UAS-nlsYFP progeny: upper dsred filter showing eye and neuronal cord fluorescence from Gal4 driver marker gene, middle cfp filter eye and neuronal cord fluorescence from UAS construct marker gene, lower YFP filter showing only background auto fluorescence from food seen also in upper and middle images.
