## Supplementary material for "Development of a functional genetic tool for Anopheles gambiae oenocyte characterisation: application to cuticular hydrocarbon synthesis": CYPG16 and CYP4G17 larval expression

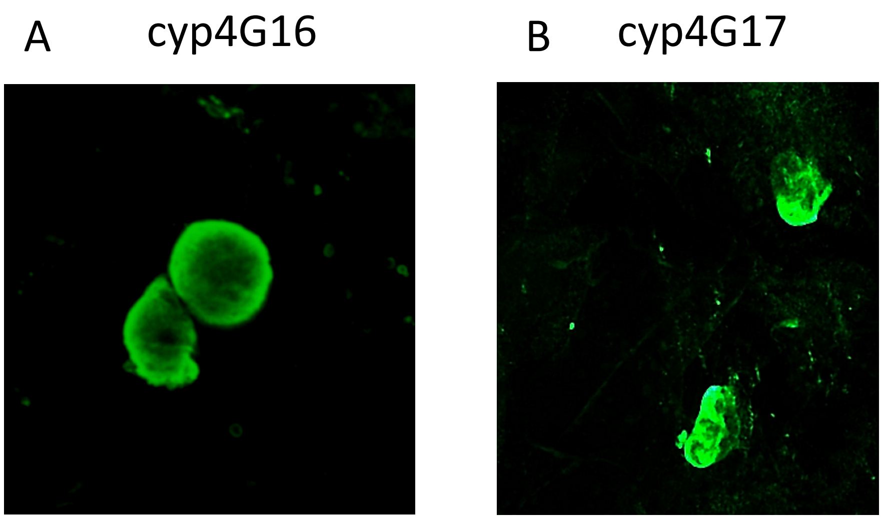


**Figure S6 CYPG16 and CYP4G17 larval expression**

Whole mount immuno histochemical stain of $4^{th}$ instar dissected abdomens with A: antisera to Cyp4g16 and B: antisera to Cyp4g17. Antisera was detected with Alexa488 (green) tagged secondary antibodies. Staining is observed in the characteristic large oenocyte cells lying at the lateral dorso-ventral junction of the larval integument.
