## Supplementary material for "Development of a functional genetic tool for Anopheles gambiae oenocyte characterisation: application to cuticular hydrocarbon synthesis": Schematic of the constructs used for gene knockout

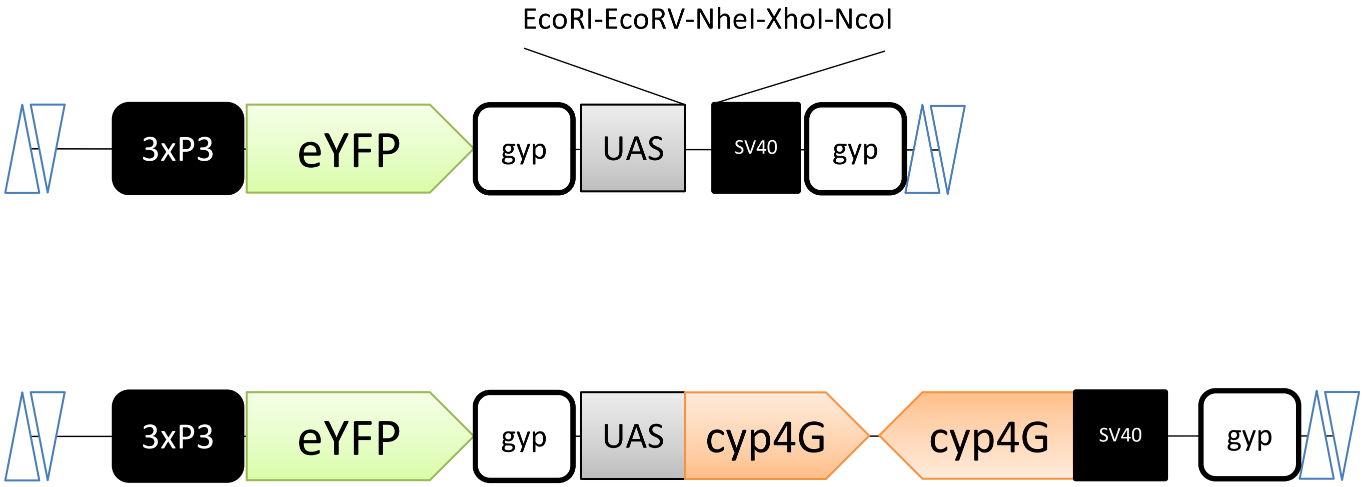


**Figure S7 Schematic of the constructs used for gene knockdown**

Knockdown plasmids based on pSL-attB-UAS-attB construct (upper). Inverted repeats of the respective *Cyp4g* fragments were inserted into the multiple cloning site of pSL-attB-UAS-attB to generate hairpin constructs (lower) described in Materials and Methods.
