## Supplementary material for "Development of a functional genetic tool for Anopheles gambiae oenocyte characterisation: application to cuticular hydrocarbon synthesis": Western analysis of 16i,17i and A14A1 early pupae

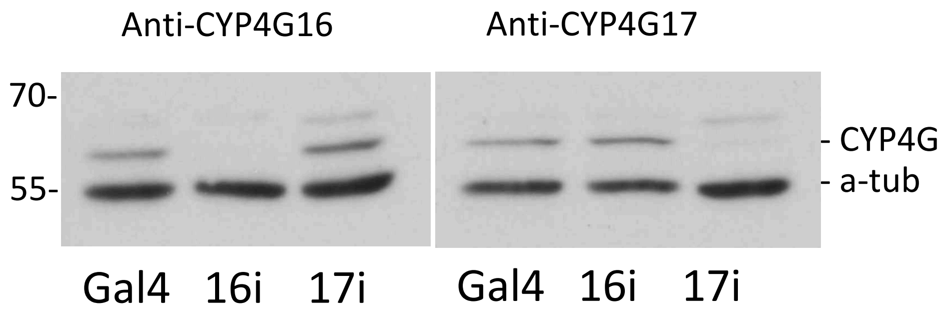


**Figure S8 Western analysis of 16i,17i and A14A1 early pupae.**

16i, 17i and sibling A14Gal4 (Gal4) extracts were blotted and screened with either CYP4G16 or CYP4G17 antisera to examine the extent of off target knockdown in either protein. Mouse anti-tubulin (a-tub) was used as an internal loading control.
