## Supplementary material for "Development of a functional genetic tool for Anopheles gambiae oenocyte characterisation: application to cuticular hydrocarbon synthesis": Plots of relative abundance of CHCs with respect to total CHC

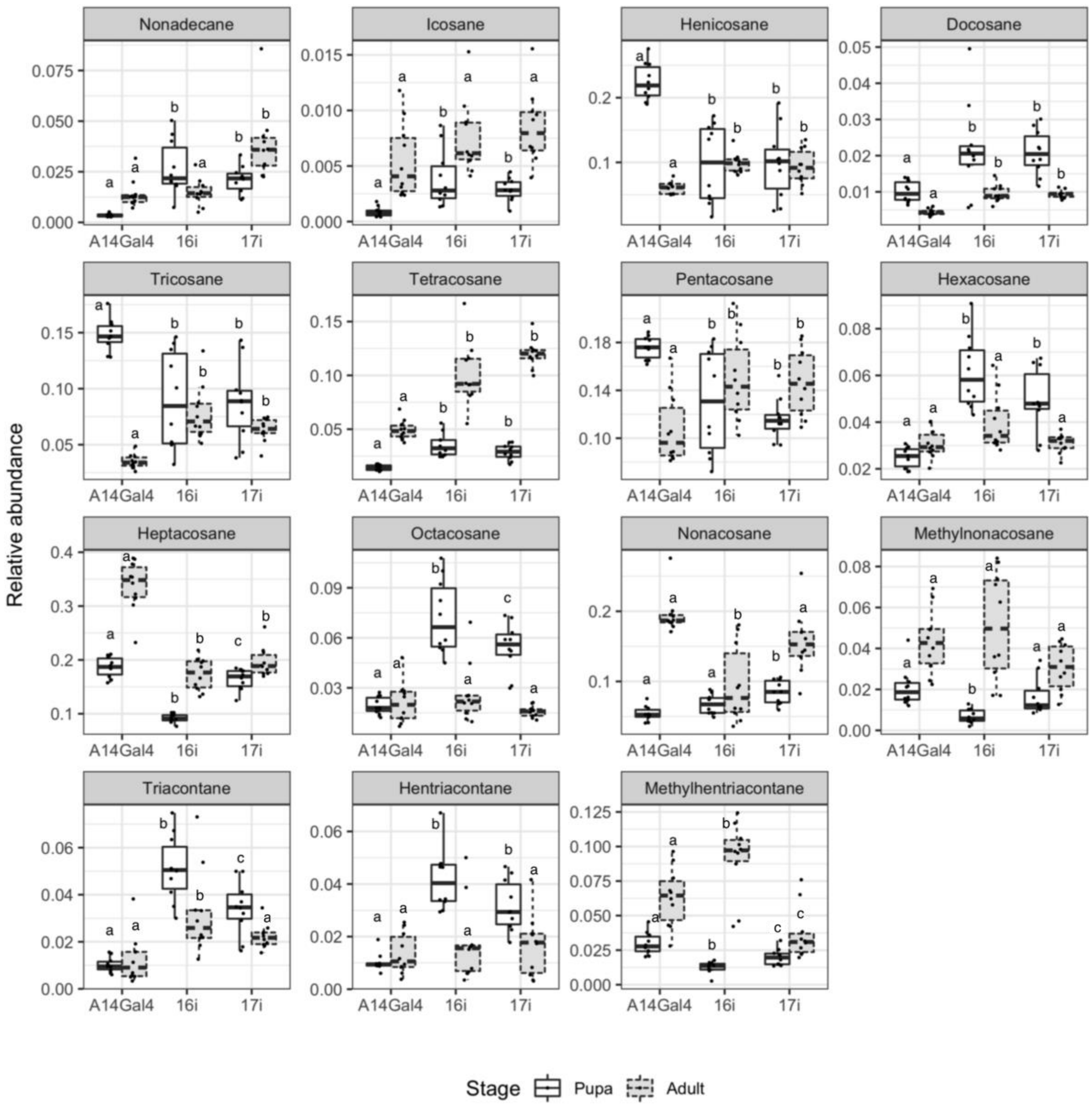


Figure S10. Plots of relative abundance of CHCs from 10 female pupae and adults. Relative abundance calculated by dividing individual peak areas by the sum of peak areas. Statistical analysis performed within stages and not across stages. Those marked with different letters are significantly different ($p<0.05$). Precise statistics given in File S3.
